# Atomic-Level Free Energy Landscape Reveals Cooperative Symport Mechanism of Melibiose Transporter

**DOI:** 10.1101/2024.08.21.608993

**Authors:** Ruibin Liang, Lan Guan

## Abstract

The Major Facilitator Superfamily (MFS) transporters are an essential class of secondary active transporters involved in various physiological and pathological processes. The melibiose permease (MelB), which catalyzes the stoichiometric symport of the disaccharide melibiose and monovalent cations (e.g., Na^+^, H^+^, or Li^+^), is a key model for understanding the cation-coupled symport mechanisms. Extensive experimental data has established that positive cooperativity between the cargo melibiose and the coupling cation is central to the symport mechanism. However, the structural and energetic origins of this cooperativity remain unclear at the atomistic level for MelB and most other coupled transporters. Here, leveraging recently resolved structures in inward- and outward-facing conformations, we employed the string method and replica-exchange umbrella sampling simulation techniques to comprehensively map the all-atom free energy landscapes of the Na^+^-coupled melibiose translocation across the MelB in *Salmonella enterica* serovar Typhimurium (MelB_St_), in comparison with the facilitated melibiose transport in a uniporter mutant. The simulation results unravel asymmetrical free energy profiles of melibiose translocation, which is tightly coupled to protein conformational changes in both the N- and C-terminal domains. Notably, the cytoplasmic release of the melibiose induces the simultaneous opening of an inner gate, resulting in a high-energy state of the system. Periplasmic sugar binding and cytoplasmic melibiose released are dynamically coupled with changes in the internal gating elements along the translocation pathway. The outward-facing sugar-bound state is thermodynamically most stable, while the occluded state is a transient state. The binding of Na^+^ facilitates melibiose translocation by increasing the melibiose-binding affinity and decreasing the overall free energy barrier and change. The cooperative binding of the two substrates results from the allosteric coupling between their binding sites instead of direct electrostatic interaction. These findings add substantial new atomic-level details into how Na+ binding facilitates melibiose translocation and deepen the fundamental understanding of the molecular basis underlying the symport mechanism of cation-coupled transporters.

## Introduction

Major facilitator superfamily (MFS) membrane transporters (MFS) are ubiquitous across all kingdoms of life, accounting for more than 25% of all transmembrane proteins, and exhibit a wide range of functions and substrate specificities. They are critical in various physiological processes, including the uptake of nutrients and drugs, as well as the expulsion of xenobiotics^1^. Certain human homologs have become promising targets for drug development due to their vital roles in nutrient and drug transport^2^. For example, the MelB homolog MFSD2A is crucial for the uptake of the essential lipid lysophosphatidylcholine in the brain^3^. Melibiose permease of *Salmonella enterica* serovar Typhimurium (MelB_St_) is a MFS sugar symporter that facilitates the simultaneous translocation of one galactoside-containing disaccharide (e.g., melibiose) and one cation (e.g., Na^+^, H^+^, or Li^+^) across the membrane in the same direction, with a strict 1:1 stoichiometric ratio ^4^ (**Fig. 1**). This symporter is a well-established and useful model for studying the cation-coupled transport mechanisms of MFS transporters^1, 4–8^. It has been experimentally well-characterized by various biochemical/biophysical techniques in combination with genetic modification^9–15^ and structural analysis^4, 16–19^. Additionally, extensive experimental data has been made available for MelB of *E. coli*^20–27^. Consistent results on the molecular recognitions for the galactoside and cation and their cooperative binding led to a hypothesis that the cooperative binding of melibiose and the coupling cation is critical for the symport mechanism^10, 28^. More recently, the correlations between substrate-binding affinities and protein conformational states have been identified through structural and binding analyses^19, 29^. While the cation-binding affinity appears largely independent of protein conformation, it was hypothesized that the sugar-binding affinity significantly decreases when MelB_St_ is in the inward-facing (IF) state^19, 29^.

**Figure 1.**
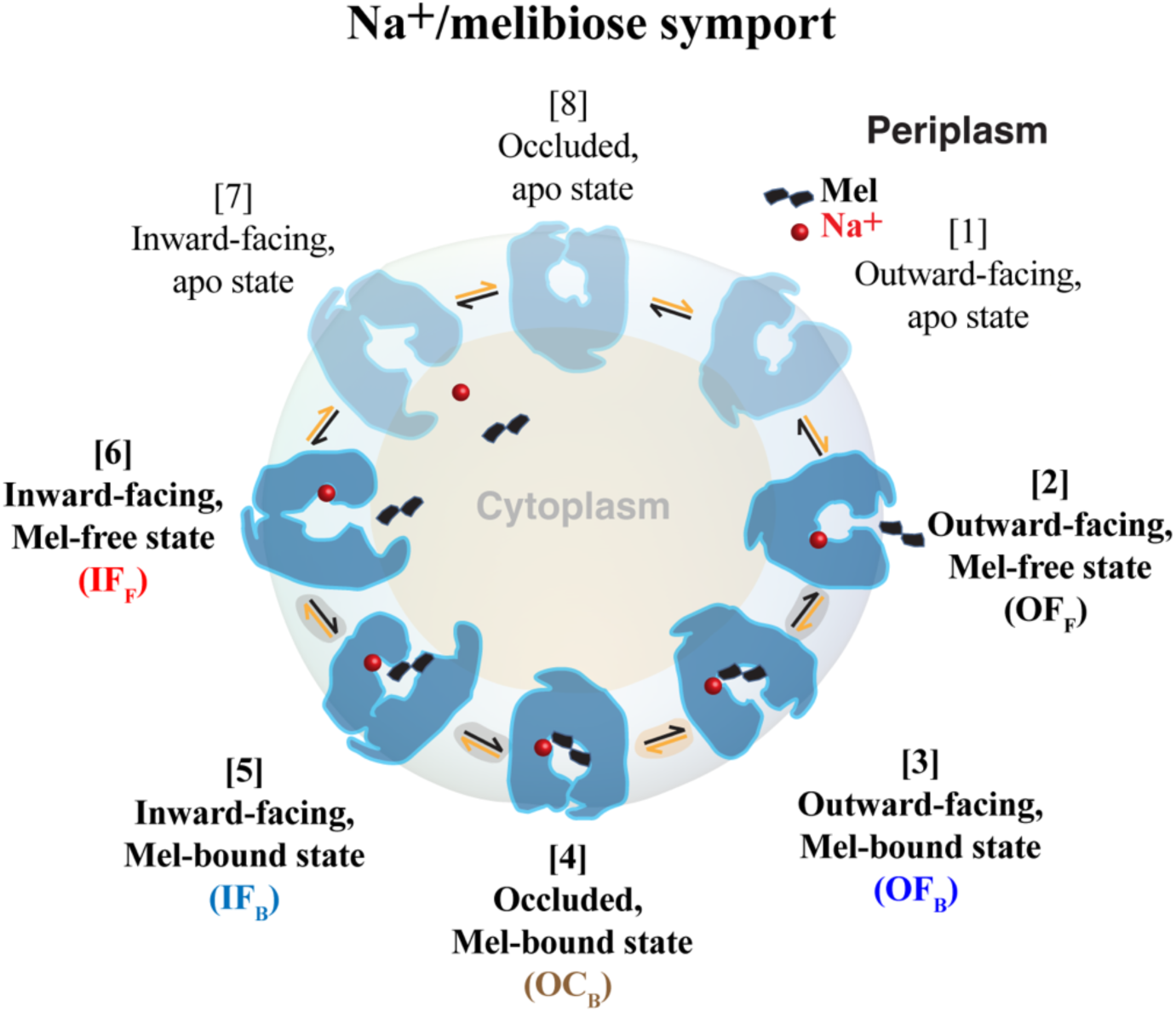
The transport cycle of the MelB symporter. The current study describes the melibiose (Mel, black double square) translocation between the periplasm and cytoplasm in the presence or absence of a bound coupling cation (Na^+^, red dot). The translocation of melibiose into cytoplasm starts from the OF_F_ state ([2]) and proceeds to the IF_F_ state ([6]) through several intermediate states such as the OF_B_ ([3]), OC_B_ ([4]), and IF_B_ ([5]). Reversal of this process results in melibiose translocation into periplasm. These key intermediate states are highlighted in bold text, with the protein depicted in solid colors. See the main text for a detailed definition of the states.

To date, the structures for two major conformational states of MelB_St_ have been resolved: the apo- or sugar-bound outward-facing (OF) state^16, 17^, and the sugar-released, Na^+^-bound IF state^18^. However, the molecular mechanisms underlying the coupled symport of the two substrates remain poorly understood. Crucial information regarding the structures, energetics and dynamics of the intermediate states (**Fig. 1**) during the transitions between the two major conformational states remains scarce. Furthermore, previous experiments have established that the binding of the two substrates is cooperative^28^, but the structural and energetic basis of this cooperativity remains unclear. Although the regions of the mobile energy barriers for regulating the sugar translocation have been identified^19^, the global free energy landscape of the entire sugar translocation process— encompassing sugar-binding and releasing events along with conformational transitions between the OF and IF conformations—remains largely unknown. The lack of quantitative characterization of such free energy landscapes has impeded a fundamental understanding of the complex interplays between the shifting of the mobile barriers and the cooperative binding of the two substrates.

Molecular simulations can fully elucidate the structures, dynamics, and thermodynamics of the transport cycle at atomic-level detail. However, calculating the sugar-translocation free energy landscape is a highly challenging task, because it requires sampling the large structural changes in the transporter coupled with substrate binding/unbinding processes. Although numerous computational studies have mapped the free energy landscapes of secondary active transporters and uniporters^30–42^, none characterized the entire substrate translocation process coupled to global protein conformational transitions in cation-coupled MFS transporters. In addition, to the best of our knowledge, there has been no study on the structural and energetic origins of the cooperative transport of two substrates in cation-coupled MFS transporters.

To this end, we calculated the free energy profiles for the translocation of the melibiose across the WT MelB_St_ in the Na^+^-bound and -unbound states, as well as the uniport D59C mutant. Building upon the recently resolved structures for both OF and IF states^17, 19^, the string method^43, 44^ was employed to identify the minimum free energy pathway (MFEP) for the translocation of the melibiose molecule from the periplasmic to cytoplasmic sides, accompanied by the OF to IF conformational transition. Extensive replica-exchange umbrella sampling (REUS) simulations were performed along the MFEP to quantify the free energy surfaces of melibiose translocation with or without a bound Na^+^. The simulations correspond to the experimentally observed melibiose exchange^4, 9, 16, 17, 45^ driven by melibiose concentration gradients without cation transduction (**Fig. 1**). Explicit characterization of the thermodynamics underlying the entire melibiose translocation process reveals the structural and energetic underpinnings of how the sugar translocation across MelB_St_ is facilitated by the binding of the coupling cation Na^+^. To the best of our knowledge, this work is the first time that the free energy landscape dictating the functional cycle of a cation-coupled MFS symporter is characterized at full atomic-level detail, thus significantly deepening our understanding of cation-coupled transporters in general.

## Results

### Melibiose translocation coupled to protein conformational changes

The **Fig. 2** depicts the free-energy profile (or potential of mean force, (PMF)) for the melibiose translocation from the periplasmic to the cytoplasmic sides of the membrane through the WT MelB_St_ with a Na^+^ bound at the cation-binding pocket. The free energy changes (*ΔG*′s) and barriers (*ΔG*^‡^′s) of key steps during the translocation are summarized in **Table 1**. The translocation process starts with a melibiose molecule in the periplasmic bulk and a MelB_St_ in its outward-facing sugar-free state (OF_F_) (image ID 0). Binding of melibiose from the periplasmic side leads to the outward-facing sugar-bound state (OF_B_) (image ID 13, **Fig. S1**), which exhibited a *ΔG* of −5.0 kcal/mol (**Table 1**, ΔG*_OF binding_*). This corresponds to a 5.0 kcal/mol melibiose binding affinity when the Na^+^-bound protein is in the OF conformation. Notably, this state has the lowest free energy in the entire translocation process and thus is thermodynamically most stable. Then, the protein conformational changes took the system from the OF_B_ to the inward-facing sugar-bound state (IF_B_) (image ID 19), overcoming a barrier of ∼6.8 kcal/mol (**Table 1**, ΔG^‡^*_OF to IF_*) required for passing through the occluded transition state (OC_B_) during the conformational transition. The OC_B_ state is a transient state, and the IF_B_ state is higher in free energy than the OF_B_ state by ∼6.4 kcal/mol (**Fig. 2**, **Table 1**). Then, the melibiose was released to the cytoplasmic side of the bulk solution, reaching the inward-facing sugar-free state (IF_B_, image ID 32), leaving behind a Na^+^-bound, sugar-free MelB_St_. This cytoplasmic melibiose release process only needs to overcome a small *ΔG*^‡^of ∼2.8 kcal/mol (**Table 1**, ΔG^‡^*_OF to IF_*) and has a small *ΔG* of ∼2.3 kcal/mol, indicating a lower melibiose-binding affinity of the IF than the OF conformations (∼5.0 kcal/mol). The entire transport process has a net *ΔG* of ∼3.7 kcal/mol (**Table 1 and Fig. 2**) and an overall *ΔG*^‡^ of ∼8.4 kcal/mol (**Table 1 and Fig. 2**).

**Figure 2.**
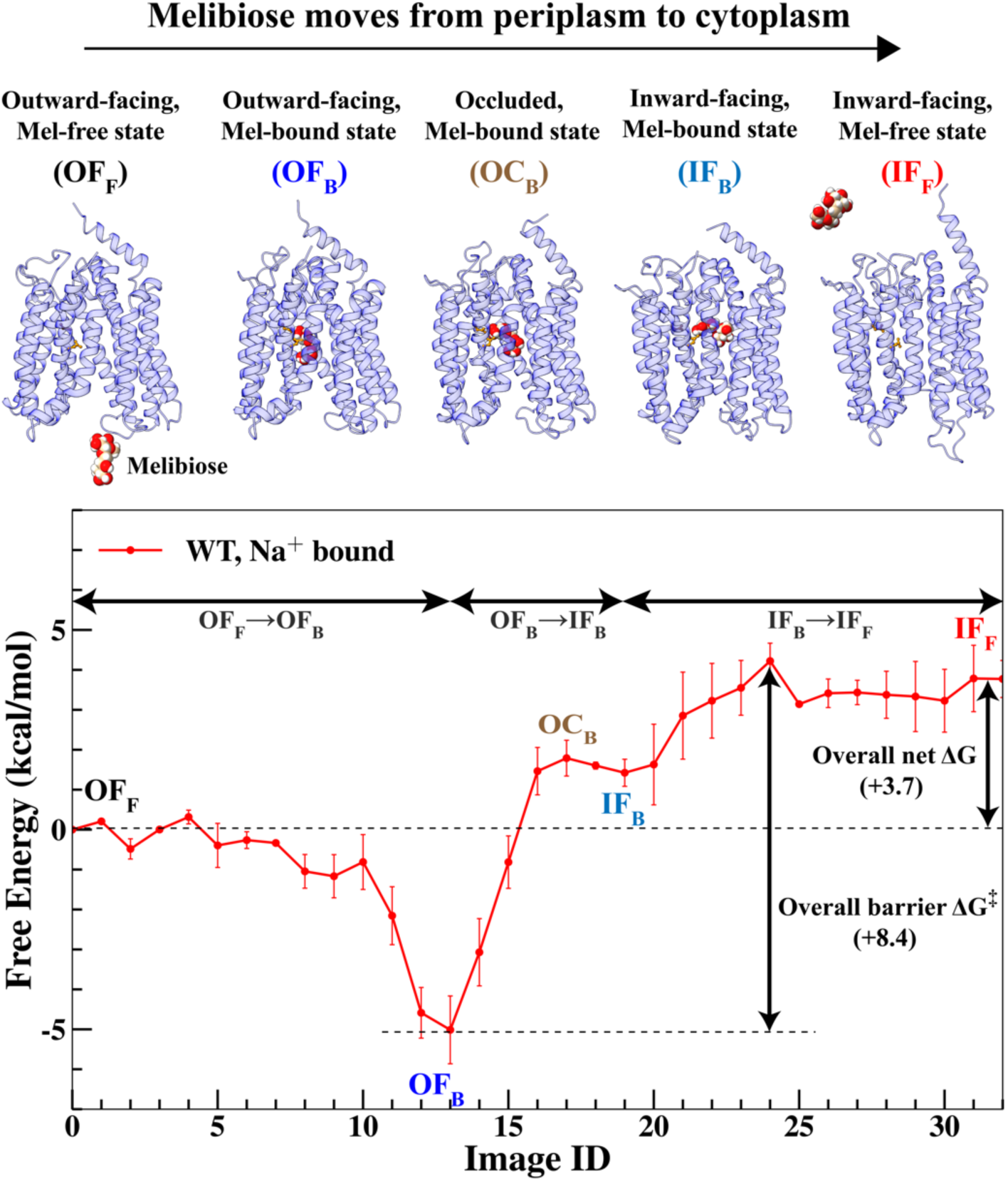
The potential of mean force (PMF) for the translocation of the melibiose across the WT MelB_St_ in the Na^+^-bound state along the minimum free energy pathway (represented as a string of images 0 to 32). Top insets: representative snapshots of the MelB_St_ (blue ribbons) and melibiose (spheres) in varied intermediate states. The bottom and top sides of each protein structure indicate the periplasmic and cytoplasmic sides of the membrane, respectively. The Asp19 and Asp124 residues, which are the major melibiose binding residues, are depicted as orange balls and sticks. The atoms in melibiose molecule are depicted as spheres.

**Table 1.**
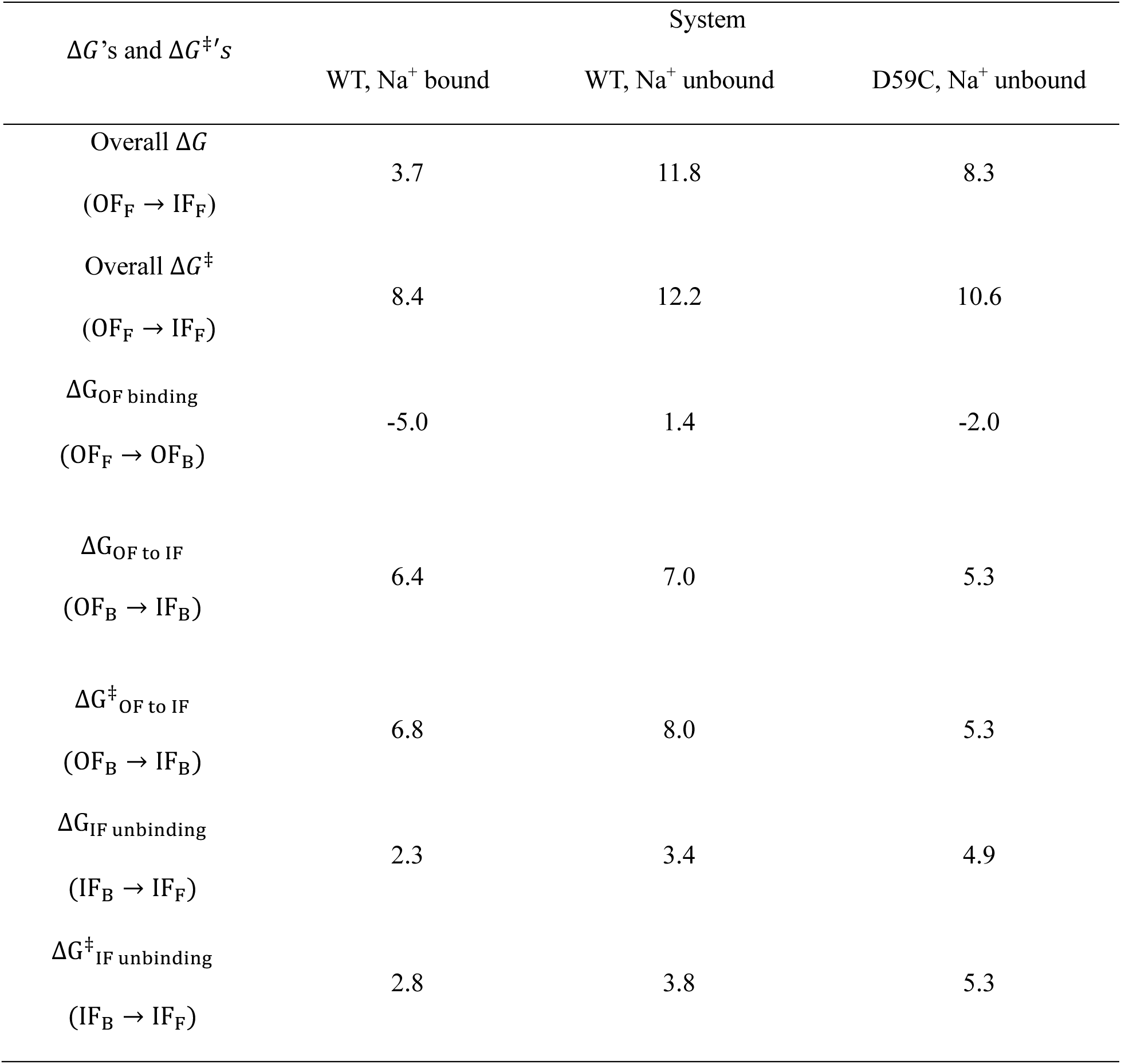
Free energy barriers (ΔG′s) and net free energy changes (ΔG^‡^′s). Free energy barriers and net free energy changes (in kcal/mol) for melibiose translocation in the WT Na^+^ bound, WT Na^+^ unbound, and the D59C mutant Na^+^ unbound systems were analyzed for the overall translocation process and several key steps in this process.

| $\Delta G$'s and $\Delta G^\ddagger$'s | System | | |
| --- | --- | --- | --- |
|  | WT, Na<sup>+</sup> bound | WT, Na<sup>+</sup> unbound | D59C, Na<sup>+</sup> unbound |
| Overall $\Delta G$<br>(OF<sub>F</sub> → IF<sub>F</sub>) | 3.7 | 11.8 | 8.3 |
| Overall $\Delta G^\ddagger$<br>(OF<sub>F</sub> → IF<sub>F</sub>) | 8.4 | 12.2 | 10.6 |
| $\Delta G_{\text{OF binding}}$<br>(OF<sub>F</sub> → OF<sub>B</sub>) | -5.0 | 1.4 | -2.0 |
| $\Delta G_{\text{OF to IF}}$<br>(OF<sub>B</sub> → IF<sub>B</sub>) | 6.4 | 7.0 | 5.3 |
| $\Delta G^\ddagger_{\text{OF to IF}}$<br>(OF<sub>B</sub> → IF<sub>B</sub>) | 6.8 | 8.0 | 5.3 |
| $\Delta G_{\text{IF unbinding}}$<br>(IF<sub>B</sub> → IF<sub>F</sub>) | 2.3 | 3.4 | 4.9 |
| $\Delta G^\ddagger_{\text{IF unbinding}}$<br>(IF<sub>B</sub> → IF<sub>F</sub>) | 2.8 | 3.8 | 5.3 |

To characterize the coupling between the melibiose translocations and protein conformational transitions, the free energy landscape was projected onto 2D free energy surfaces (FES) spanned by different combinations of collective variables (CVs) using Eq S4 (**Fig. 3**). In **Fig. 3A**, the 2D FES is spanned by the first principle component (PC1) of the backbone Cα atoms of all 12 transmembrane helices and the melibiose transport coordinate Z (see Method in SI for definition). As Z increases from −30 Å to +30 Å, the melibiose molecule moves through MelB_St_ from the periplasmic to the cytoplasmic sides of the membrane. As the PC1 gradually increases from ∼-3.5 Å to +3 Å, the transmembrane helices transit from the OF to the IF conformational states.

**Figure 3.**
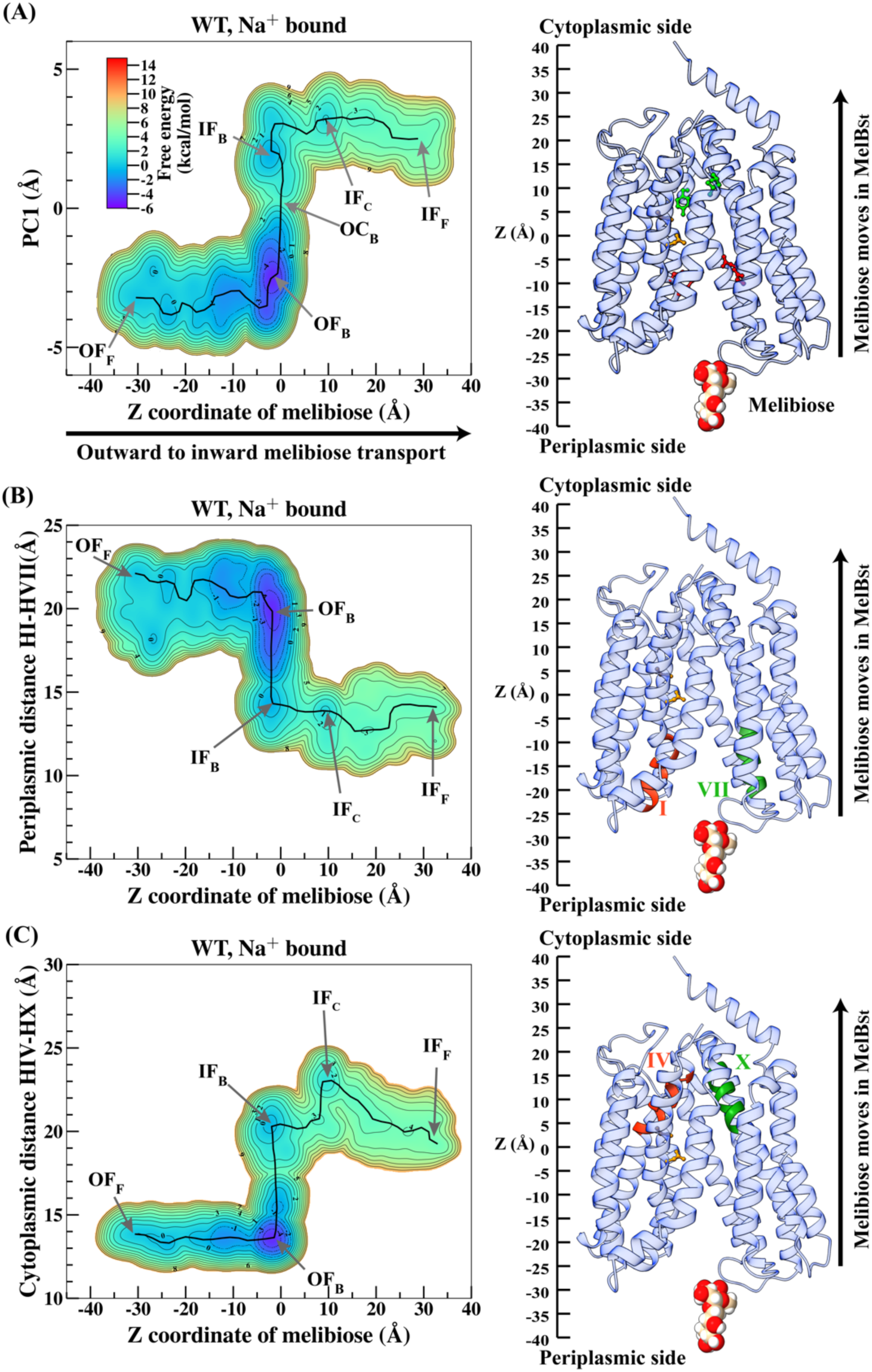
Free energy surfaces (FES) for the transport of the melibiose through the WT MelB_St_ in the Na^+^ bound state. (A) FES spanned by the first principle component of the backbone (PC1) vs. melibiose transport coordinate Z. An illustrative structure indicating the scale of the Z coordinate is shown on the right, and key residues in the periplasmic gate, binding site and cytoplasmic gate are highlighted in red, orange and green, respectively (see main text). (B) FES spanned by interhelical distance between helices I and VII on the periplasmic side. An illustrative structure indicating the helices I (red) and VII (green) is shown on the right. (C) FES spanned by interhelical distance between helices IV and X on the cytoplasmic side. An illustrative structure indicating the helices IV (red) and X (green) is shown on the right. The approximate minimum free energy pathways (MFEP) tracking the major basins on the FES are indicated as black lines, and the regions corresponding to the intermediate states are labeled by arrows. All FES plots share the same color bar as (A).

As shown in **Fig. 3A**, the periplasmic sugar-binding process is correlated with dynamic fluctuations of the transmembrane helices, as exhibited by the shifts in the PC1 values across the multiple minima encountered in the OF_F_→OF_B_ process. For example, from OF_F_ (Z = −30 Å, PC1 = −3.2 Å) to OF_B_ (Z = −3 Å, PC1 = −2.5 Å), the PC1 values fluctuate back and forth and has a net increase of ∼0.7 Å after reaching the OF_B_ minimum. These fluctuations in PC1 correspond to non-negligible conformational fluctuations in the overall transmembrane protein backbones during the periplasmic-binding process. The binding-induced conformational change is more evident when the FES is projected to the 2D plane spanned by the translocation coordinate and the interhelical distance between helix I and VII near the periplasmic side of the membrane (**Fig. 3B**). There is a decrease in this interhelical distance from ∼22 Å at OF_F_ to ∼19 Å at OF_B_ as the melibiose approaches the binding site. Thus, the periplasmic sugar-binding slightly closes the periplasmic side of the sugar translocation pathway, preparing the protein for the subsequent conformational changes.

After the periplasmic sugar-binding event, the OF_B_→IF_B_ process begins, largely increasing the PC1 from approximately −2.5 Å to +2 Å (**Fig. 3A**). This process features global protein conformational changes critical for the entire transport process. In this step, the sugar translocation pathway is fully closed on the periplasmic side and opened on the cytoplasmic side, preparing the protein for releasing sugar into the cytoplasm. A large decrease in multiple periplasmic interhelical distances is observed, such as between helices I and VII (from ∼19 Å to ∼14 Å) (**Fig. 3B**), as well as an increase in cytoplasmic interhelical distances, such as between the helices IV and X (from ∼13 Å to ∼20 Å) (**Fig. 3C**).

After the system reaches the IF_B_ state, the melibiose release to the cytoplasmic bulk begins (IF_B_⟶IF_F_). The process goes through an intermediate state IF_C_, where the melibiose passes through the most constricted region of the entire pathway contributed by Val145, Val346 (helix X), and Tyr369 (helix XI) (**Fig. 3A**). From IF_B_ to IF_C_, the cytoplasmic inter-helices distances further increase. For example, the distance between helices IV and X further increases to ∼23 Å near Z = + 11 Å to prepare the space for the melibiose to pass through (**Fig. 3A**). After this, the melibiose is eventually released to the cytoplasmic bulk and the interhelical distance between helices IV and X slightly decreases back to below 20 Å. The diagonal nature of the pathway linking IF_B_, IF_C_, and IF_F_ (**Fig. 3C**) indicates that the motion of the substrate and the expansion/shrinkage of the cytoplasmic constricted region are tightly coupled. The finding is significant because it reveals that bound melibiose does not passively wait for the cytoplasmic path to widen, as previously speculated. Instead, it actively induces protein conformational changes to move out. This observation redefines the alternation-access model, demonstrating that both the periplasmic binding and cytoplasmic release of melibiose are tightly coupled with protein conformational changes.

### Changes in the pore radius profile during melibiose translocation

The coupling between the conformational change and the melibiose translocation is also evident through analyzing the radius profiles of the internal cavities that form the sugar translocation path in different intermediate states. For example, we first monitored the radius of the cavity near the periplasmic bulk (Z = −25 Å) during the entire sugar translocation process. This radius slightly decreases during periplasmic binding (OF_F_→OF_B_, red vs. orange curves in **Fig. 4A**). It then largely decreases by more than 3 Å during conformational transition (OF_B_→IF_B_, orange vs. green curves in **Fig. 4A**). Following this, it remains almost the same during cytoplasmic release (IF_B_→IF_F_, green vs. blue curves in **Fig. 4A**). Monitoring the radius of the cavity near the cytoplasmic bulk (Z = + 25 Å), it remains small below 2 Å during the periplasmic binding, and then increases by approximately 3 Å following the OF_B_→IF_B_ transition. Thus, in general, the OF_F_→IF_F_ process is accompanied by the widening and shrinking of the periplasmic and cytoplasmic terminals of the path in a reciprocal manner.

**Figure 4.**
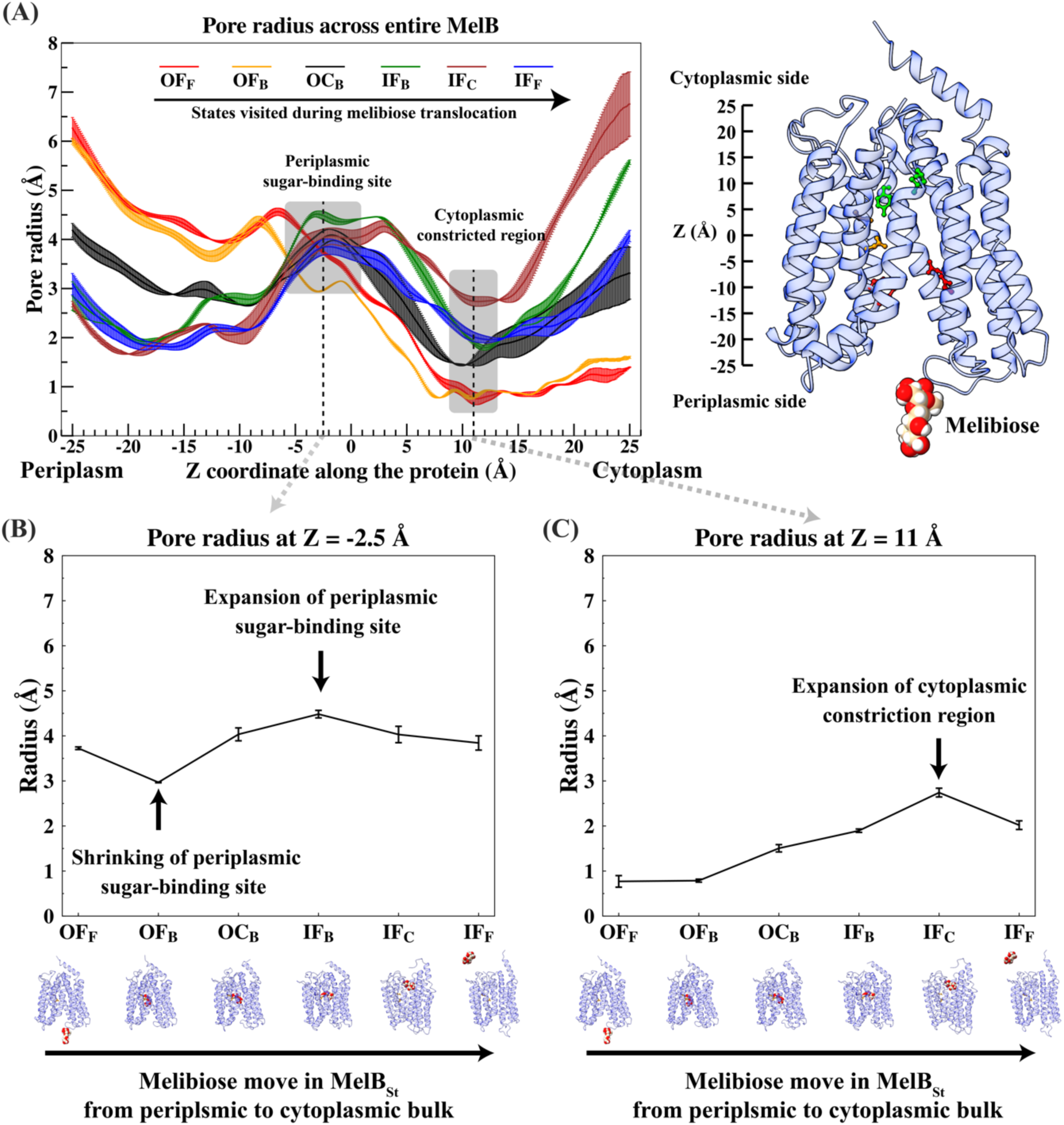
Change of pore radius profile coupled to melibiose transport through WT MelB_St_ in the Na^+^ bound state. (A): Pore radii profile across the MelB as a function of the Z coordinate (relative to the COM of Asp124 and Asp19) along the entire protein (x-axis) for each different state during the melibiose translocation. As the melibiose is translocated from the periplasm to cytoplasm, the system transitions from the OF_F_ (red) to IF_F_ (blue) states through the OF_B_ (orange), OC_B_ (black), IF_B_ (green), and IF_C_ (brown) intermediate states, each of which features different pore radius profiles. The IF_C_ state corresponds to the melibiose passing through the cytoplasmic constricted region near Z=11 Å. An illustrative structure indicating the scale of the Z coordinate is shown on the right. (B) and (C): Translocation of melibiose from the periplasmic to cytoplasmic sides of MelB_St_ induces the change of pore radii measured at Z=-2.5 Å (periplasmic sugar-binding site) and Z=11 Å (cytoplasmic constricted region). The melibiose translocation process is represented as the transitioning of the system from the OF_F_ to IF_F_ states through multiple intermediate states along the x-axes. The protein structures below the plots serve as visual guides for the location of the melibiose in different states.

The radius profile also revealed two constricted regions surrounding the bound sugar molecular, which functions as gates on both the periplasmic and cytoplasmic sides of the sugar translocation path. The periplasmic constriction region is located near Z = −15 to −10 Å and formed by the cavity-lining residues, particularly the Tyr26 and Met27 on the kink of the helix I, and Asn248 and Asn251 on the helix VII (**Fig. 4A, right panel**). Clearly, the helix I kink moves towards the helix VII positioning in the middle of the melibiose-accessing path upon melibiose binding. Tyr26 frequently interacts with the melibiose as it passes through and defines the periplasmic edge of the sugar-binding pocket, and its extended bulky sidechain partially occludes the bound sugar from the periplasmic side. This constricted region is further narrowed when MelB_St_ changes to the occluded transition state. Thus, the Tyr26 functions as a key gating residue. Experimentally, mutation of the Tyr26 to Cys26 residue resulted in the loss of active transport of the MelB_St_, highlighting the importance of this residue in the functional cycle^13^. The cytoplasmic constriction region is located near Z = +11 Å, which is formed by the cavity-lining residues Val145, Val346 and Tyr369, as discussed above. When the periplasmic side of the transporter is open, the N- and C-terminal domains on the cytoplasmic side form a tightly connected salt-bridge network, as described previously^13, 17^. These ionic interactions majorly contribute to forming the inner barrier of cytoplasmic sugar release. As the melibiose passes through this region, the radius in this region expands to accommodate its motion (**Fig. 4C**), in line with the above-mentioned observation from the FES (**Fig. 3C**).

Another important observation is that periplasmic binding reduces the radius of the sugar-binding pocket (**Fig. 4B**), reaffirming the binding-induced conformational change discussed above (**Fig. 3**). The subsequent OF to IF conformational changes (OF_B_ →IF_B_) increase the radius of the sugar-binding cavity near Z = −2.5 Å (**Fig. 4B**). The expansion of the binding pocket eventually facilitates the release of melibiose to the cytoplasmic side. This is consistent with the results from the free energy profile calculation. The simulation results thus corroborate the previous experimental result^19^ that the outward-facing conformation has a higher sugar-binding affinity than the inward-facing (**Fig. 2-3**) and explain its structural and energetic origin with atomic-level detail.

### Cooperative motion of N- and C-terminal domains

Multiple helices in both the N- and C-terminal domains change their relative orientation with respect to the normal of the membrane plane during the melibiose translocation. To quantify this observation, the free energy of the system is projected to 2D surfaces spanned by two collective variables (CVs), which measure the directional tilt angle of a helix with respect to the membrane plane normal (see “Method” for definition) and melibiose translocation. Based on the topographies of the 2D FES’s (**Fig. S4 B-G**), it is obvious that in the N-terminal domain, all helices decrease their tilting angles by 10-15 degrees as Z moves from −35 to 35 Å. In contrast, in the C-terminal domain, all helices increase their tilt angles by 10-20 degrees as Z moves from −35 to 35 Å (**Fig. S4 H-M**). The simultaneous tilting of all helices suggests that both domains must reorient with respect to the membrane normal to complete the global protein conformational transition. This transition exhibits a symmetric-like movement of the two pseudo-symmetric helix bundles. Notably, this finding, which cannot be derived from experimental structures alone due to the lack of a membrane surface, helps resolve a longstanding debate regarding the alternating-access model.

### Melibiose translocation energetically coupled to Na^+^ binding

Next, we characterize the PMF of melibiose translocation when the MelB_St_ is in the Na^+^-unbound state (**Fig. 5**, blue curve; **Table 1**). The comparison between the PMFs of Na^+^-unbound and the Na^+^-bound states is essential for understanding how the Na^+^ binding cooperates with the thermodynamics and kinetics of melibiose translocation. When the Na^+^ is unbound, the overall net *ΔG* and overall barrier *ΔG*^‡^are ∼11.8 kcal/mol and ∼12.2 kcal/mol, respectively (**Table 1**). These values are greater than the Na^+^-bound state by 8.1 kcal/mol and 3.8 kcal/mol, respectively, indicating that the melibiose translocation is thermodynamically and kinetically less favorable if the Na^+^ is unbound. Several factors contribute to the more endergonic and slower translocation process in the absence of bound Na^+^. First, periplasmic melibiose binding (image IDs from 0 to 15) is endergonic with ΔG*_OF binding_* ∼ 1.4 kcal/mol, in contrast to the exergonic process in the Na^+^ bound state (ΔG*_OF binding_* ∼ −5.0 kcal/mol). Second, the endergonic protein conformational transition (image IDs from 15 to 21) exhibits ΔG*_OF to IF_* ∼ 7.0 kcal/mol and ΔG^‡^*_OF to IF_* ∼ 8.0 kcal/mol, which are both greater than the Na^+^ bound state (6.4 kcal/mol and 6.8 kcal/mol, respectively). Third, the endergonic cytoplasmic unbinding event (images 21 to 32) exhibits ΔG*_IF binding_* ∼ 3.4 kcal/mol and ΔG^‡^*_IF unbinding_* ∼ 3.8 kcal/mol, again greater than the Na^+^ bound state (2.3 kcal/mol and 2.8 kcal/mol, respectively).

**Figure 5.**
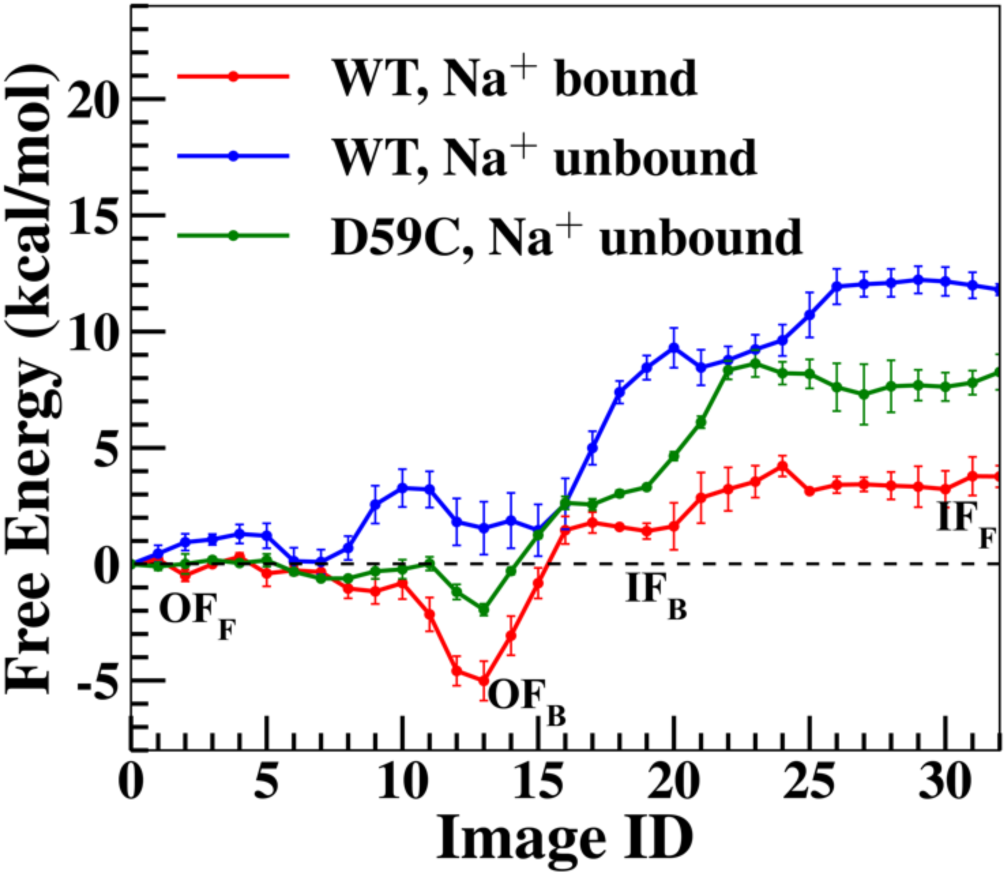
PMFs for the transport of the melibiose across MelB_St_ along the MFEPs represented as strings of images (0 to 33). The PMFs are calculated in the WT, Na^+^ bound (red) and unbound (blue) states, and the D59C mutant in the Na^+^ unbound state (green).

The well-characterized D59C mutation results in a melibiose uniporter due to the loss of the carboxyl group on the Asp59, which is the only residue critical for binding of all three types of coupled cations (Na^+^, H^+^, and Li^+^)^12, 18, 28^. This mutant thus loses cation-coupled active transport activity but can facilitate the transport of melibiose downhill its concentration gradient^17^ and the exchange of melibiose across the membrane ^16^. This unique feature makes the mutant a good model system for understanding the Na^+^/melibiose symport mechanism.

The melibiose translocation across the D59C mutant has an overall net *ΔG* of ∼ 8.3 kcal/mol and overall free energy barrier *ΔG*^‡^ of 10.6 kcal/mol (**Fig. 5**, green curve; **Table 1**). They are both greater than the WT Na^+^ bound system (3.7 kcal/mol and 8.4 kcal/mol, respectively), making the melibiose translocation thermodynamically and kinetically less favorable. Several factors contribute to the more endergonic and slower melibiose translocation in the mutant. The periplasmic sugar binding (image IDs from 0 to 13) has a ΔG*_OF binding_* of ∼ −2.0 kcal/mol, indicating that the mutation diminishes the binding affinity of the OF state by ∼3.0 kcal/mol as compared to the WT Na^+^-bound system. Unlike the WT Na^+^-unbound system, the periplasmic binding in the D59C mutant is exergonic despite its incapability of binding cations, which is consistent with previous experiential measurements^17^. After the OF_B_ state is reached, the OF_B_⟶IF_B_ transition together with the cytoplasmic sugar release (IF_B_⟶IF_F_) (image IDs from 13 to 32) results in a combined high barrier of ∼10.6 kcal/mol (ΔG*_OF to IF_* + ΔG^‡^*_IF unbinding_*). This rate-limiting barrier in the mutant is ∼2.2 kcal/mol higher than the WT Na^+^ bound system. Notably, the mutant features ∼3 kcal/mol higher barrier for the cytoplasmic sugar release, which is likely caused by the different orientations of the residues lining the cytoplasmic constricted region.

In both Na^+^ unbound systems (WT and D59C), as expected, melibiose translocation is still coupled with protein conformational changes, but to a lesser extent than in the WT, Na^+^ bound system (**Figs. S5-S10**). In both Na^+^ unbound systems, periplasmic binding and cytoplasmic release are coupled with changes in the overall conformation of the protein and specific interhelical distances (**Figs. S5-S6**), as well as pore radius of the sugar-binding pocket and cytoplasmic constricted region (**Figs. S7-S8**). These changes are smaller in scale compared to the WT though. The directional tilt angles of helices in the N- and C-terminal domains follow the same trend as the WT Na^+^ bound system (**Figs. S9-S10**).

Overall, the removal of Na^+^ from the cation-binding site, either due to low Na^+^ concentrations in bulk or loss of Na^+^ binding site, can largely reduce melibiose binding affinity, as has been well-documented by a variety of experimental tests in varied methods^9, 12, 16–18, 28^. This is one of the key factors contributing to the increased overall *ΔG*’s and *ΔG*^‡^’s for the melibiose translocation in the two Na^+^ unbound systems. These results explicitly demonstrate that the free energy profiles explicitly demonstrate that the co-transport of Na^+^ and melibiose is a direct result of binding cooperativity between the two substrates.

### Allosteric coupling between Na^+^ and melibiose binding

To understand the different melibiose binding affinities in the WT Na^+^ bound and unbound systems, we performed a total of 500 ns unbiased MD simulations in the OF_B_ state for each system. The interaction energies between the melibiose and MelB_St_ are compared between these two systems (**Fig. S11)**. The interaction energy is decomposed into contributions from electrostatic and van der Waals components (**Fig. S11 A&B,** green and red bars, respectively). Importantly, the direct interaction energy between the bound Na^+^ and melibiose in the WT Na^+^ bound system was also evaluated.

Upon the unbinding of Na^+^ (**Fig. S11A)**, the protein-melibiose interaction energy shifted from −113 ± 2 to −105.4 ± 0.5 kcal/mol in WT MelB_St_, consistent with the reduction in the binding affinity of the melibiose. Such a reduction largely arises from the electrostatic component, which changes from −96 ± 2 to −88.0 ± 0.4 kcal/mol. Notably, the direct interaction energy between the bound Na^+^ and melibiose (∼ 0.2 ± 0.5 kcal/mol) in the WT Na^+^ bound state is negligible. This finding is important in that the decrease in the melibiose binding affinity upon Na^+^ unbinding is not due to losing the direct Na^+^-melibiose interaction but due to a weakened protein-melibiose interaction. The decreased melibiose-binding affinity in the absence of Na^+^ is mainly attributed to the alteration of local electrostatic interactions between melibiose and its nearby residues (**Fig. S11B)**.

The WT Na^+^ bound system features a narrower distribution of the local melibiose-protein interaction energy that peaked at a lower energy range than the WT Na^+^ unbound system (**Fig. S12A)**. This implies that the Na^+^ unbinding alters the binding cavity and disfavors melibiose binding. To further probe the structural origin of this effect, we analyzed the hydrogen-bonding interactions between the melibiose and protein (**Table S1)**. Na^+^ unbinding from the WT MelB_St_ led to the loss of 0.9 ± 0.2 hydrogen bonds between the melibiose and protein. In addition, the hydrogen bonds with three charged residues Asp124, Asp19, and Arg149, which contribute most to the protein-melibiose hydrogen bonds in the OF_B_ state, are affected by the Na^+^ unbinding the most. Thus, the reduction of hydrogen bonds between melibiose with the protein contributes to decreased binding affinity in the absence of a bound Na^+^ ion.

The change in the hydrogen-bonding interaction due to Na^+^ unbinding likely results from the allosteric coupling between the cation-binding and melibiose-binding sites. The Asp124 and Tyr120 residues in helix IV are essential in the coupling pathway since this helix also contributes the Thr121 residue to the Na^+^-binding site. Without a bound Na^+^ ion, the distance between the Asp55 and Asp124 is elongated (**Fig. S12B**). The same trend is also observed for the distances between Asp124 and Asp19 (**Fig. S12C)**, between Tyr120 and Asp55 (**Fig. S12D**), and between Tyr120 and Asp19 (**Fig. S12E**). Furthermore, the distance between Asp55 and Asp19, i.e., the two key residues in the cation- and sugar-binding sites, respectively, is increased upon Na^+^ unbinding (**Fig. S12F**). The correlations between the changes in these pairwise distances suggest that both Asp124 and Tyr120 residues propagate structural changes from the Na^+^-binding site to the sugar-binding site upon Na^+^ unbinding, leading to a decrease in melibiose affinity. The unbinding of Na^+^ weakens the hydrogen bond between Asp19 and the 3-hydroxyl group of melibiose, as indicated by their elongated separation distance (**Fig. S12G**). This observation aligns with previous experimental studies^13, 16, 24, 27, 46^: the D124C or Y120C mutants can bind melibiose but lose Na^+^-coupled melibiose active transport^16^. Our simulations thus provide an atomistic-level explanation for these experimental observations in these decoupling mutants.

## Discussion

Calculating the free energy landscape that describes the protein conformational transitions associated with substrate translocation through transporters is a challenging yet essential task for understanding their functional mechanisms. In this study, building upon two resolved ligand-bound structures representing the inward-^19^ and outward-facing conformational states^17, 18^, we uncovered a wealth of new mechanistic information about the functional cycle of MelB. This was achieved through extensive free energy calculations combined with state-of-the-art reaction path-finding techniques. Notably, the use of two experimental structures as initial guesses for the two endpoints of the minimum free energy pathway (outward-facing sugar-free and inward-facing sugar-free states) enhances the reliability of the string method (**Fig. S3**). This is an improvement over previous applications of this approach to other transporters, where only one major conformational state (outward-facing, inward-facing, or occluded) was experimentally available, and the other major conformations had to be generated by biased molecular dynamics simulations^31, 33, 35^. Our simulation results are consistent with a large body of experimental data collected in the past decades via varied biochemical and biophysical data and structural analyses^9, 10, 12, 13, 18, 19, 28^.

First, all simulation data consistently showed that Na^+^ binding increases the melibiose-binding affinity of WT MelB_St_, in agreement with previous experimental measurements^9, 28^. Second, the free-energy landscape indicates that the inward-facing states are thermodynamically less stable than the outward-facing states, regardless of the binding of Na^+^ and melibiose, providing the energetic data supporting the conclusion drawn from structural analysis^19^. Third, the overall free energy landscape reveals that the outward-facing conformation of the WT has the highest sugar-binding affinity, and the OF_B_ state is the thermodynamically most stable state for the entire translocation process (**Table 1 & Fig. 2**). This is a piece of direct evidence supporting that the IF conformational state has a lower sugar-binding affinity and the experimentally measured binding affinity is primarily related to the OF conformational state^19^.

During the melibiose translocation, only a modest rate-limiting barrier is encountered in the OF_B_→OF_F_ process (∼5 kcal/mol, **Fig. 2**). The protein conformational transition and cytoplasmic release is a cooperative process that increases the free energy of the system. The opening of the cytoplasmic gate is narrow and is induced by the motion of the sugar, so the cytoplasmic sugar-release process can be appropriately described as sugar squeezing through the inner barrier. In addition, the lack of a thermodynamically stable occluded state (OC_B_) facilitates the OF_B_→IF_B_ transition. All these features make MelB_St_ a highly effective transporter.

The free energy landscapes reveal the coupling mechanisms between the two substrates and the transporter, providing new insights into experimental measurements of MelB_St_ and its mutants^10, 17, 19, 28^. Without Na^+^, both WT and D59C mutants exhibit higher *ΔG*^‡^and *ΔG* than the WT Na^+^-bound system, making melibiose translocation less favorable. This suggests that the coupling between sugar and cation is primarily due to energetic coupling at the binding step. Na^+^<u>binding increases melibiose affinity, lowers the free energy barrier and change, and facilitates the protein conformational transition, thus accelerating melibiose translocation.</u> The molecular basis underlying the difference in the free energy landscapes involves strengthened hydrogen bonds between melibiose and its binding residues (Asp19, Asp124, Arg149) due to Na^+^ binding at the cation-binding pocket (Asp55, Asp59, Thr121, Asn58), with allosteric coupling through Asp124 and Tyr120 on helix IV, as proposed sed on experimental data^16, 24, 27, 46^. The interaction between the two binding sites underpins the energetic and kinetic coupling necessary for the cooperative transport of Na^+^ and melibiose.

The WT MelB_St_ mediates both the active transport and facilitated diffusion of melibiose ^4^. In contrast, the D59C mutation eliminates the active transport mode of WT but retains its melibiose uniport activity^16, 17^, similar to the E325A mutant of LacY^7, 47^. Although such a phenomenon is observed in the D59C MelB and has been an important clue for identifying the cation-binding site of cation-coupled secondary transporters (7, 47, 48), the molecular origin of why the mutant retains uniport activity remains elusive to date. Here, the comparison between the free energy landscapes of the WT and mutant offers a microscopic-level explanation. For the D59C mutant, the overall *ΔG*^‡^ of melibiose translocation is modestly higher than WT, Na+ bound MelB_St_ by only 2.2 kcal/mol (Table 1). Thus, this mutation does not kinetically block the inward sugar translocation entirely, and it can even facilitate the outward translocation process by reducing the periplasmic sugar-releasing barrier as a result of reduced binding affinity in the OF state (**Fig. 5**). Therefore, the bi-directional exchange and gradient-downhill transport of melibiose can still be observed so long as the melibiose concentration is high enough on at least one side of the membrane. Notably, the coupling between the melibiose translocation and conformational change in this uniporter mutant remains (Figs. S6, S8 & S10). From an evolutional point of view, it is likely that the introduction of a cation site into a uniporter can accelerate the substrate translocation and increase the substrate-binding affinity. This symport function evolved from a uniporter can better serve cellular needs by harvesting substrate from scarce conditions.

In summary, our all-atom free energy landscapes of MelB_St_ provided numerous critical, novel and fundamental insights into the structural and energetic origins of the coupling mechanism essential in cation-coupled MFS symporters: (1) the substrate translocation is tightly coupled to global protein conformational changes where all transmembrane helices are reoriented with respect to the membrane normal, (2) periplasmic binding is coupled with a partial closure of the periplasmic gate and shrinkage of sugar-binding site, (3) the outward-facing sugar-bound state is thermodynamically most stable during the entire translocation process, (4) the occluded state is a transient state, (5) the melibiose cytoplasmic release is coupled to the temporary expansion of the cytoplasmic gate, and (6) the allosteric coupling between the cation- and sugar-binding sites leads to cooperative binding of Na^+^ and melibiose, which facilitates sugar translocation thermodynamically and kinetically. For the first time, our free energy profiles explicitly demonstrate that the co-transport of Na^+^ and melibiose is a direct result of binding cooperativity between the two substrates. Notably, although such cooperativity has been observed experimentally on a macroscopic level, our free-energy simulations comprehensively characterized this core mechanism of symport with atomic-level detail, and provide explicit, crucial evidence for energetic coupling between the two co-transported solutes across all intermediate states visited during the entire melibiose translocation process (**Figs. 1-2 & 5**).

## Methods

A brief summary of the methods is provided here, with full details included in the SI.

### System setup

Three simulation systems were prepared to calculate the free energy landscapes of melibiose translocation coupled with protein conformational transition: (1) WT MelB_St_ with Na^+^ bound, (2) WT MelB_St_ without Na^+^, and (3) D59C mutant without Na^+^. The OF and IF conformational states were derived from crystal (PDB code 7L16^17^) and CryoEM (PDB code 8T60^19^) structures, respectively. Key steps in the structural preparation included mutating residues to match the desired states, positioning melibiose in the binding site, adding a lipid bilayer, and neutralizing with 0.15M NaCl. Each system consisted of ∼130,000 atoms in a simulation box of ∼100 ×100 ×130 Å³. All systems were constructed using the CharmmGUI web interface^48^.

### Preparation for string method simulations

Initial string images for string method simulations were prepared for all three systems. First, for melibiose binding/unbinding processes in OF and IF states, the system was relaxed with geometry optimization and then equilibrated with backbone restraints. The bound melibiose was gradually pulled out of the protein (cytoplasmic unbinding for the IF state and periplasmic unbinding for the OF state) using a series of harmonic potentials, creating intermediate snapshots for substrate release into both sides of the membrane. This provided initial string images for OF_F_→OF_B_ and IF_B_→IF_F_ transitions. Next, the geodesic interpolation algorithm^49^ generated 10 intermediate images of the transmembrane backbone for the OF_B_→IF_B_ conformational transition. These backbone structures guided restrained simulations that dynamically relaxed all atoms in the images. All simulations used the CHARMM36m^50–56^ and TIP3P force fields^57^, and were performed using the NAMD software package ^58^.

### String method simulations

The String Method with Swarms of Trajectories (SMwST)^43, 44^ was employed to identify the minimum free energy pathway (MFEP) for melibiose translocation in all three systems. This method has been successfully applied to other types of secondary active transporters^31, 33, 35^ and protein complexes^59^. The initial string consisted of 33 images, representing the entire OF_F_ ↔ IF_F_ transition. These images were projected into a 13-dimensional space defined by CVs related to melibiose translocation and protein conformational changes.

The SMwST simulations involved 500 iterations, where each image was equilibrated and propagated in each iteration. The string was updated iteratively until convergence was achieved, as monitored by RMSD in the 13-D space **(Fig. S2)**. The endpoints of the final string corresponded closely to the experimental OF and IF structures **(Fig. S3)**.

### Replica-exchange umbrella sampling (REUS) simulations

The last iteration of the string from the SMwST simulation was used as the MFEP for performing REUS simulations^60, 61^. The 33 image centers were interpolated into equidistant 13-D window centers for 33 umbrella windows. A harmonic potential (0.5-1 kcal/mol) acted on each of the 13 CVs per window. Initial conditions for each window were taken from the images in the last SMwST iteration. Replica exchanges among 10 neighboring windows were attempted every 10 ps on a rotation basis^59^. The REUS simulations ran for 90 ns per window, with the first 10 ns discarded as equilibration, resulting in 7.9 μs of total sampling time for all three systems.

### Analysis

The REUS simulations were unbiased using a generalized version^33^ of the weighted histogram analysis method^62, 63^ (Eqs. S1-S2). The free energies of each image were corrected using Eq S3. The 2D free energy surfaces were constructed using Eq. S4. The MFEP connecting the OF_F_ and IF_F_ states on the 2D free energy surfaces (FES) was traced using the zero-temperature string method. The HOLE program^64^ was used to analyze the pore radius profiles for all snapshots sampled by the REUS simulations.

## Author contributions

The manuscript was written through the contributions of all authors. Ruibin Liang designed the research, performed the simulations, analyzed the data, and wrote the manuscript. Lan Guan designed the research and revised the manuscript. All authors have given approval to the final version of the manuscript.

## Data Availability Statement

The data supporting this article have been included in the Supplementary Information.

## Supplemental information

The supplemental information contains **Figure S1-S12**. It includes a snapshot of the melibiose binding site in the WT, Na^+^ bound MelB_St_, the evolution of RMSD of the string in the 13-dimensional CV space as a function of iteration number during the string method simulation for all three systems, the backbone RMSD of the string images with respect to the experimental structures, the free energy surfaces and pore radius profiles during the entire melibiose translocation across the WT and D59C mutant of MelB_St_ in the Na^+^ unbound state, the energy decomposition analysis of Na^+^-protein interaction, and the distances between the coupling residues (D124 and Y120) to the cation-binding and sugar-binding sites. The supplemental information also contains the MFEPs of all three systems as separate text files.

This information is available online, free of charge.

## Supporting information

Converged string images

## Acknowledgments

This work was supported by the National Institutes of Health Grants R35GM150780 to R.L. and R35GM153222 to L.G. The researchers used GPU computing facilities provided by the High-Performance Computing Center at Texas Tech University. The authors also acknowledge the helpful insights and inputs provided by Prof. Abhishek Singharoy at Arizona State University, and the initial simulation system setup by Mr. Amirhossein Bakhtiiari at Texas Tech University.

---

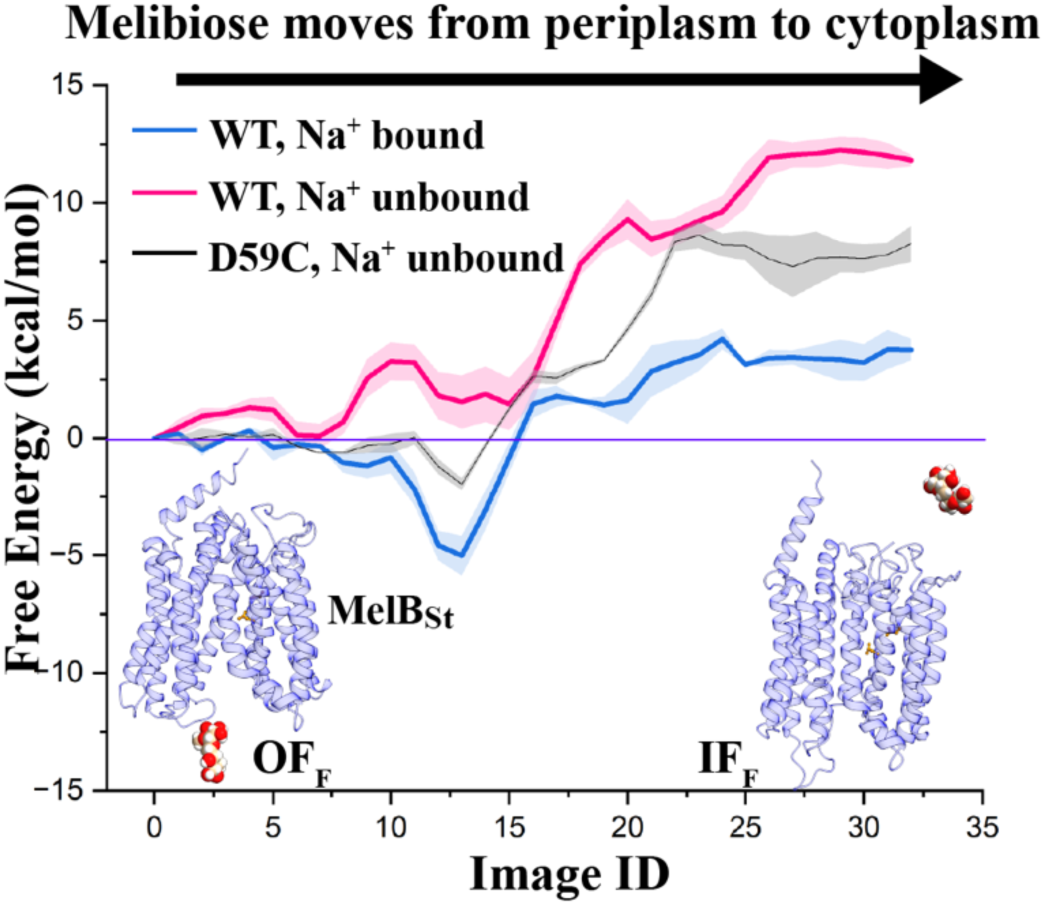

All-atom free energy landscapes of the entire melibiose translocation process in wild-type MelB_St_ and its uniport D59C mutant elucidate the structural and energetic basis of the positive cooperativity between melibiose and its driving cation. The new insights significantly deepen our understanding of the molecular basis underlying cation-coupled transport mechanisms in MFS symporters.

