## Supplementary material for "Atomic-Level Free Energy Landscape Reveals Cooperative Symport Mechanism of Melibiose Transporter": SI text and figures

### Supplemental Information for Atomic-Level Free Energy Landscape Reveals Cooperative Symport Mechanism of Melibiose Transporter

*Ruibin Liang[1,3*] and Lan Guan[2*]*

1. Department of Chemistry and Biochemistry, Texas Tech University, Lubbock, United States
2. Department of Cell Physiology and Molecular Biophysics, Center for Membrane Protein Research, Texas Tech University Health Sciences Center, School of Medicine, Lubbock, United States
3. Lead contact

AUTHOR INFORMATION

*** Corresponding authors**

#### Method

##### *System setup*

In total, three simulation systems were set up in preparation for calculating the free energy landscapes of melibiose translocation coupled with protein conformational transition: (1) wild-type (WT), $Na^+$ bound state of $MelB_{St}$, (2) WT, $Na^+$ unbound state of $MelB_{St}$ and (3) the D59C mutant, $Na^+$ unbound state of $MelB_{St}$. For each system, the OF and IF conformational states were built from a previously resolved crystal structure (PDB code 7L16)[1] and a CryoEM (PDB code 8T60)[2] structure, respectively. The OF structure is for the D59C mutant [1], while the IF structure is for WT $MelB_{St}$ complexed with a nanobody [2]. All organic solvents were removed. The nanobody was removed from the IF structure. The Cys59 residue in the OF structure was mutated back to Asp59 to generate the initial OF structure for the two WT $MelB_{St}$ systems. The Asp59 residue in the IF structure was mutated to Cys59 to generate the initial IF structure for the D59C mutant. A melibiose molecule was positioned in the carbohydrate-binding site by keeping only the disaccharide moiety of the detergent ligand in the OF crystal structure. For the IF structure, the following procedure was employed to generate the initial coordinates of the melibiose molecule. First, the OF system (with melibiose) was aligned with the IF CryoEM structure using the transmembrane protein backbone heavy atoms as the reference atoms for minimizing the root mean square displacements (RMSD). Then, the melibiose's coordinates in the aligned OF system were directly used as the initial coordinates of melibiose in the IF system, whose protein-heavy atom coordinates were obtained from the CryoEM structure. Then, the hydrogen atoms were added to the protein-heavy atoms following standard protonation states (deprotonated Asp and Glu, neutral His with a proton on the delta nitrogen, protonated Ser, Thr, Cys, Tyr, Arg, and Lys). A mixed lipid bilayer with a total of 288 lipid molecules was added to embed the protein in the XY-plane, following a POPE:POPG ratio of 7:2 to mimic the membrane composition of *E. coli*. For the WT, $Na^+$ bound state, a sodium ion was manually inserted in the binding site surrounded by the side chains of Asp55, Asp59, Asn58, and Thr121, following our previous studies[3, 4]. For the $Na^+$-unbound state of both WT and D59C mutant, the $Na^+$ binding site was left empty. Then, the membrane-embedded MelB was capped by two boxes of water molecules with 25 Å thickness, one above and the other below the lipid bilayer. Each system was neutralized with 0.15M NaCl to mimic physiological conditions. The setups resulted in periodic boundary condition (PBC) simulation boxes of ~$100 \times 100 \times 130$ Å$^3$, consisting of ~130,000 atoms. The CharmmGUI web interface [5] was utilized to construct the system.

##### *Preparation for string method simulations*

The following procedures were followed to prepare for the initial string images of string method simulations for all three systems. Firstly, the images for the melibiose binding/unbinding processes in the OF and IF conformational states of MelB (i.e., $OF_F \leftrightarrow OF_B$ and $IF_B \leftrightarrow IF_F$ in **Fig. 2**) were prepared. Initially, 1000 steps of geometry optimization were used to relax the melibiose-bound IF and OF systems ($IF_B$ and $OF_B$) with harmonic restraints (500 kJ/mol/Å$^2$ force constant) applied to the heavy atoms of the protein and lipids. Subsequently, a 250 ps simulation at a temperature of

303.15 K was conducted in the constant NVT ensemble. The temperature of the system was maintained using a Langevin thermostat with a friction coefficient of 1 $ps^{-1}$. The restraints were then systematically diminished to 0.1 kcal/mol during 2 ns of dynamics in the constant NPT ensemble at 303.15 K and 1 atm pressure. The pressure of the system was maintained using the Langevin piston Nose-Hoover method.[6, 7] Then, constant NPT ensemble simulations at the same temperature and pressure were performed for 10 ns, with a week restraint of 5 kcal/mol/$Å^2$ acting on the RMSD of transmembrane protein $C_\alpha$ atoms with respect to the experimental structures (PDB codes 7L16 and 8T60 for the OF and IF systems, respectively) [1, 2]. Such constraints were found to be necessary to temporarily keep the melibiose-bound MelB structures close to the experimental ones, because melibiose binding is coupled with conformational changes, especially for the thermodynamically less stable $IF_B$ state (see **Results**).

Starting from the preliminarily equilibrated $IF_B$ and $OF_B$ structures, the bound melibiose molecule was gradually pulled out of the MelB to both the periplasmic and cytoplasmic sides of the membrane. In each direction, a 4 kcal/mol/$Å^2$ harmonic potential was imposed on the collective variable (CV), defined as the distance from the center of mass (COM) of melibiose to that of Asp19 and Asp124 residues, projected onto the Z-axis. The harmonic potential's centers were changed by 2.4 Å every 200 ps until the melibiose fully exits the MelB from either the periplasmic or cytoplasmic side of the pore (CV > 29 Å for the cytoplasmic side and CV < -32 Å for the periplasmic side). This pulling process was performed with the above-mentioned RMSD restraints on the protein $C_\alpha$ atoms to maintain the OF, and IF structures of the protein in all windows, and it created a series of umbrella windows with the melibiose restrained at different locations across the pore. Then, the system in each window was equilibrated with the corresponding harmonic restraint center and the above-mentioned force constant for another 10 ns. Importantly, during this equilibration step, no restraint on the protein was imposed, which allows the relaxation of the protein and solvent (water and lipid) in response to the different positions of the melibiose. The final coordinates and velocities of different windows in this step prepared the initial string images for the $OF_F \rightarrow OF_B$ and $IF_B \rightarrow IF_F$ processes (see below).

Secondly, the following procedure was followed to prepare the initial string images for the conformational transition from outward-facing melibiose-bound ($OF_B$) state to inward-facing melibiose-bound state ($IF_B$) (i.e., $OF_B \leftrightarrow IF_B$). First, the geodesic interpolation algorithm [8] was utilized to interpolate all protein $C_\alpha$ atoms in the equilibrated $IF_B$ and $OF_B$ structures, which generated 10 images of protein $C_\alpha$ atoms gradually transitioning from the $OF_B$ structure to $IF_B$ structure. This interpolation algorithm has been tested to generate a smooth transition pathway using an internal coordinate representation, which avoids high-energy intermediate structures [8]. Therefore, it can serve as a decent initial guess for the minimum free energy pathway (MFEP) connecting the two endpoints. Next, the 10 images of $C_\alpha$ atoms were utilized as reference structures for restraining the $OF_B$ and $IF_B$ structures to the intermediate structures during the $OF_B \leftrightarrow IF_B$ transition. For each image, the restrained simulations were performed in two stages to gradually move the protein backbone to the reference structure. First, the system was equilibrated for 100 ps with an RMSD harmonic restraint of 500 kcal/mol/Å$^2$ force constant acting on all protein $C_\alpha$ atoms. Following this, the system was equilibrated for another 1 ns with positional harmonic restraints of 1 kcal/mol/Å$^2$ acting on all protein $C_\alpha$ atoms. Simultaneously, a harmonic potential of 4 kcal/mol was added to the melibiose-pulling CV (see above) to keep the melibiose close to the equilibrated binding site (near CV = -2.5 Å) during the protein's conformational biasing process. The last snapshot from each of the 10 restrained simulations constituted the 10 images describing the $OF_B$ ↔ $IF_B$ transition in the initial guess of the string.

For all equilibration simulations, a 2 fs timestep was employed to propagate the trajectory, with constraints on bond lengths involving hydrogen atoms. The CHARMM36m force field [9-15] was employed for the protein, lipids, and melibiose, and the TIP3P model for water molecules [16]. Electrostatic interactions were computed using the particle mesh Ewald method [17] with a 12 Å cutoff for van der Waals interactions. All MD simulations were performed using the NAMD software package [18].

***String method simulations***

The string method with swarms of trajectories (SMwST) algorithm [19, 20] was utilized to identify the MFEP for melibiose translocation across the MelB in all three systems. The SMwST simulations were initiated from an initial string consisting of 33 images/ The images with IDs of 0 to 10, 11 to 21, and 22 to 32 describe the $OF_F$ ↔ $OF_B$, $OF_B$ ↔ $IF_B$ and $IF_B$ ↔ $IF_F$ processes, respectively. The coordinates and velocities of the images for the initial string were obtained from the above-mentioned procedure. In the SMwST simulations, each image was projected to a 13-dimensional space spanned by 13 CVs. The first CV was identical to the one employed for pulling the melibiose out of the MelB from the binding site, which played a major role in describing the translocation of the melibiose. The remaining 12 CVs were designed to describe the conformational transition from the OF to IF structures, as detailed below. The twelve helices of MelB surrounding the melibiose transport pore were separated into six pairs (helices I & VII, II & VIII, III & IX, IV & X, V & XI, VI & XII). Within each pair, the two helices approximately reside on the opposite side of each other across the central pore. For each pair, 2 CVs are defined, one as the $C\alpha$ atoms' COM distance between the cytoplasmic segment of the two helices (**Fig. 3 C**), and the other as the $C\alpha$ atoms' COM distance between the periplasmic segments of the helices (**Fig. 3 B)**. All distances were projected onto the XY-plane. Thus, the increase in the 6 CVs describing the interhelical distances near the cytoplasmic side and the decrease in the 6 CVs on the periplasmic side corresponds to the OF to IF conformational transition.

With the 13 CVs and initial string defined, the SMwST was performed for 500 string update iterations in the 13-D conformational subspace spanned by the CVs. In each iteration for each image, the system was first restrained and equilibrated near the 13-D image center using a harmonic restraint acting on each of the 13 CVs with a force constant of 20 kcal/mol. The equilibration run lasted for 20 ps. Then, a swarm of 50 unbiased trajectories was launched from the equilibrated structure and velocity and propagated for 40 fs while the temperature and pressure were maintained at 303.15 K and 1 atm by Langevin thermostat and Langevin piston Nose-Hoover method. Due to the stochastic dynamics, the 50 trajectories are non-identical. Afterward, the average displacement along each CV was calculated from the 50 unbiased trajectories and added to the corresponding CV of this 13-D image center. Such a procedure was repeated 33 times to update all image centers in each of the 500 string update iterations. Following this, an equidistant interpolation of all updated image centers yielded the new image centers that make up the new string, from which the next iteration of the string update began. In our test, 500 iterations were sufficient to converge to a dynamically stable string as monitored by the evolution of its RMSD in the 13-D space using the initial iteration as the reference (**Fig. S2**). The endpoints of the converged strings have backbones in the transmembrane region resembling the OF and IF experimental structures (**Fig. S3**).

***REUS simulations***

The last iteration of the string obtained from the SMwST simulation was utilized as the MFEP to perform the replica-exchange umbrella sampling (REUS) simulation[21, 22]. Specifically, the 33 image centers after the last iteration were interpolated equidistantly along the MFEP, generating 13-D window centers for 33 umbrella windows. For each window, a harmonic potential of 0.5-1 kcal/mol acted upon each of the 13 CVs defined for the SMwST simulation. The initial coordinates and velocities of the system in each window were obtained from the corresponding image in the last iteration of the SMwST simulation. Throughout the simulation, the exchanges among replicas were attempted every 10 ps. For each replica, each exchange attempt was with one of the replicas in its 10 closest neighboring windows on a rotation basis [23]. The REUS simulations were propagated for 90 ns for each of the 33 windows. The first 10 ns of the trajectory was discarded as an equilibration, and the last 80 ns was treated as the production trajectory for free energy calculation and structural analysis. The total REUS sampling time for all windows of all three systems was thus ~7.9 μs.

***Analysis***

The free energy of each umbrella window was calculated by unbiasing the CV distributions from the REUS simulations using a generalized version [24] of the weighted histogram analysis method [25, 26] (generalized WHAM). The perturbed free energy for each window ($F_i's$, up to an additive constant) and the unbiased weight for each snapshot $t$ ($\omega^t$'s) were calculated by iterating over Eq 1 & 2 until convergence:

$$e^{-\beta F_i} = \sum_t \omega^t e^{-\beta U_i(\vec{\zeta}^t)} \qquad \text{Eq S1}$$

$$\omega^t = \left(\sum_i T_i e^{-\beta\left(U_i(\vec{\zeta}^t) - F_i\right)}\right)^{-1} \qquad \text{Eq S2}$$

In Eq 1, $\vec{\zeta}^t$ is the 13-D CVs at snapshot $t$, $U_i(\vec{\zeta}^t)$ is the energy contributed by the harmonic potential of window $i$ for snapshot $t$, $\omega^t$ is the unbiased weight of snapshot $t$, and the sum on the right hand side is over all snapshots sampled by all windows. In Eq 2, $T_i$ is the number of snapshots sampled by window $i$, and the sum on the right-hand side is over all windows. The $F_i$'s were used to construct the final PMF for the melibiose transport process using Eq 3 under the stiff-spring approximation:[24]

$$G\left(\vec{\zeta}(s)\right) \approx F(s) + \frac{1}{2\beta k}\left(\beta\left(\frac{d}{ds}F(s)\right)^2 - \frac{d^2}{ds^2}F(s)\right), \qquad \text{Eq S3}$$

where the $G\left(\vec{\zeta}(s)\right)$ is the final PMF result, $s$ is the arc length along the 13-D string starting from image 0, $\vec{\zeta}(s)$ is a 13-D point at $s$, $F(s) = F(\vec{\zeta}(s))$ is the perturbed free energy at $s$ (interpolated from $F_i$'s), $\beta = 1/k_BT$, $\frac{d}{ds}F(s)$ and $\frac{d^2}{ds^2}F(s)$ are the first and second derivatives of $F(s)$ evaluated at $s$, $k$ is the force constant in the harmonic biasing potential used in the REUS. For visual comparison between PMFs under different states, $G\left(\vec{\zeta}(s)\right)$ was evaluated at all images at $s_i$'s ($0 \leq i \leq 32$) arc length along the pathway. The error bars of $G\left(\vec{\zeta}(s)\right)$ were estimated by block-average analysis.

Using the unbiased weights ($\omega^t$'s), the free energy surface (FES) in the subspace spanned by combinations of CVs ($\vec{\xi}$) other than the 13 CVs used in the REUS were reconstructed following Eq 4:

$$G(\vec{\xi}) = -\beta^{-1}\log\left(\sum_t \omega^t K\left(\vec{\xi}(\vec{x}^t) - \vec{\xi}\right)\right), \qquad \text{Eq S4}$$

where the $\vec{x}^t$ are the all-atom coordinates of snapshot $t$, $\vec{\xi}(\vec{x}^t)$ are the positions of the multidimensional CVs at snapshot $t$, $K$ is a kernel function, $\omega^t$ is the unbiased weight of snapshot $t$, and the sum in the log function is over all frames in all windows. Eq 4 was used for reconstructing the 2D FES on the plane spanned by the first principle component (PC1) of transmembrane protein C$\alpha$ atoms, and CV was used to describe the transport of the melibiose in REUS. Before the principle component analysis (PCA), the transmembrane protein C$\alpha$ atoms in all snapshots were first aligned to the average structure of all frames, and the PCA was performed on only the transmembrane protein C$\alpha$ atoms to save computational cost. Then, each snapshot's transmembrane protein C$\alpha$ coordinates were projected to the PC1 and combined with the corresponding value of the melibiose transport CV to generate the 2D $\vec{\xi}(\vec{x}^t)$. Eq 4 was also used for reconstructing 2D FES spanned by the melibiose translocation CV and other CVs describing the interhelical distances and the directional tilt angle of helices (see below).

The MFEP connecting the $OF_F$ and $IF_F$ states on some of the 2D FES's (**Fig. 3** and **Figs. S4-S5)** were traced using zero-temperature string method with 60 images. The 2D strings evolved based on the gradients of the 2D FES until convergence, after which they approximate the MFEP on these surfaces.

The HOLE program[27] was used to analyze the pore radius profile for all snapshots sampled by the REUS. The pore radius profiles for each state in each system (**Fig. 4** and **Figs. S7-8**) were calculated by weighted averaging over all snapshots in the windows corresponding to that state using the $\omega^t$ of each snapshot. The error bars of the pore radius profiles were estimated by block-average analysis.

***Definition of the directional tilt angle of helices***

A specialized CV was designed to monitor the changes in the directional tilting of the 12 transmembrane helices relative to the membrane surface normal during melibiose translocation. For any given snapshot of REUS, a 2D plane is first established using two vectors. The first vector is the positive Z-axis, approximating the average membrane normal throughout the simulation. The second vector, labeled as the "D axis" in **Fig. S4A, S9-S10**, points from the center of mass (COM) of the N-terminal domain to the COM of the C-terminal domain in that snapshot and is projected onto the XY-plane, which approximates the average membrane plane. The vector along the principal axis of a helix (extending from the periplasmic to the cytoplasmic ends) is then projected onto this 2D plane defined by the Z and D axes. The angle between the projected vector and the Z-axis is defined as the directional tilt angle. The tilt angle is defined as positive if the angle between the projected vector and the positive direction of the D axis is less than 90 degrees; otherwise, it is negative. In other words, a N-terminal domain helix has a positive tilt angle if its cytoplasmic part is closer to the C-terminal domain than its periplasmic part, and vice versa. A C-terminal domain helix has a positive tilt angle if its cytoplasmic part is farther from the N-terminal domain than its periplasmic part, and vice versa.

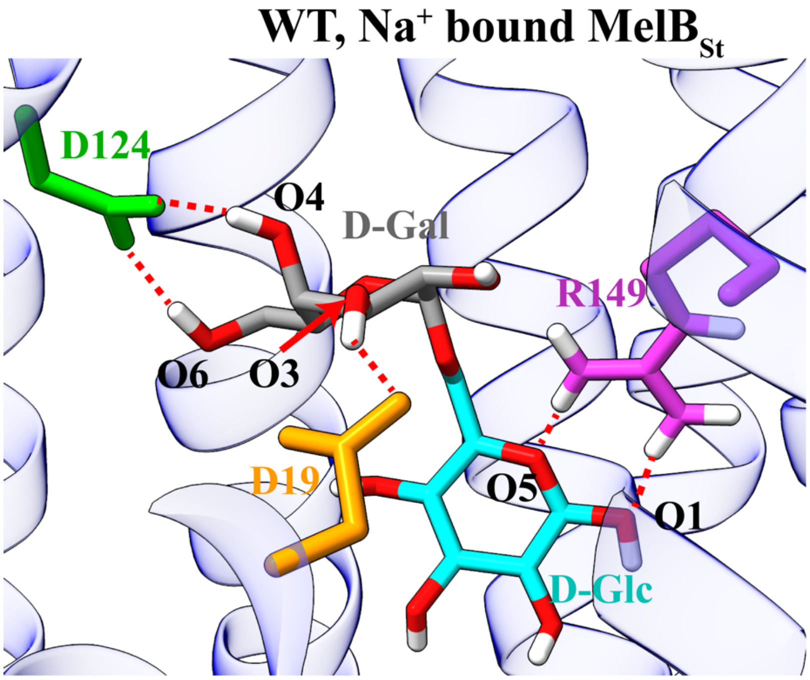


**Figure S1.** Structure of the melibiose binding site in MelBSt. The residues D124, D19, and R149, shown in green, orange, and purple respectively, form hydrogen bonds with melibiose. The galactosyl and glycosyl moieties are depicted in gray and cyan. The galactosyl moiety forms hydrogen bonds with residues D124 and D19 through the hydroxyl groups at the C3, C4, and C6 positions. These specific interactions are crucial for the selective binding of galactosides by MelB.

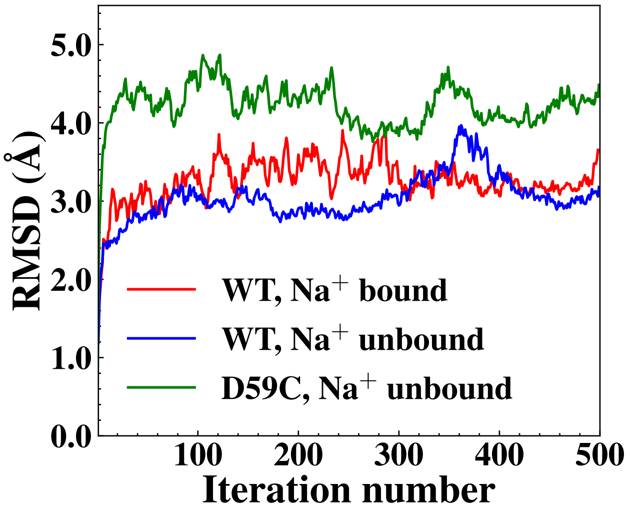


**Figure S2.** The root mean square displacements (RMSDs) of the strings in the 13-dimensional CV space with respect to the initial string as a function of iteration number in the string method simulation.

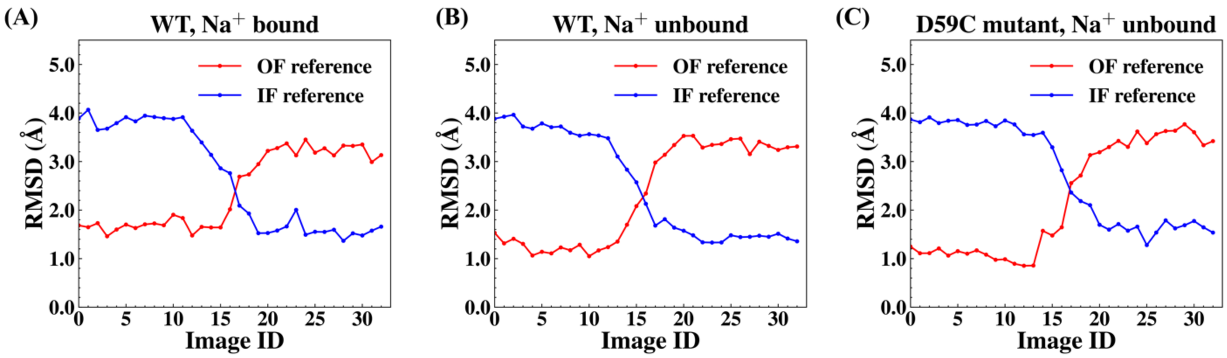


**Figure S3.** Protein backbone RMSD between the images of the converged string in each system with respect to the experimental structures in the outward-facing (PDB code 7L16) and inward-facing (PDB code 8T60) conformational states. For each image, the Cα atoms of the transmembrane region of the $MelB_{St}$ are compared with corresponding atoms in the experimental structures. (A) WT, $Na^+$ bound system. (B) WT, Na+ unbound system. (C) D59C mutant, $Na^+$ unbound system.

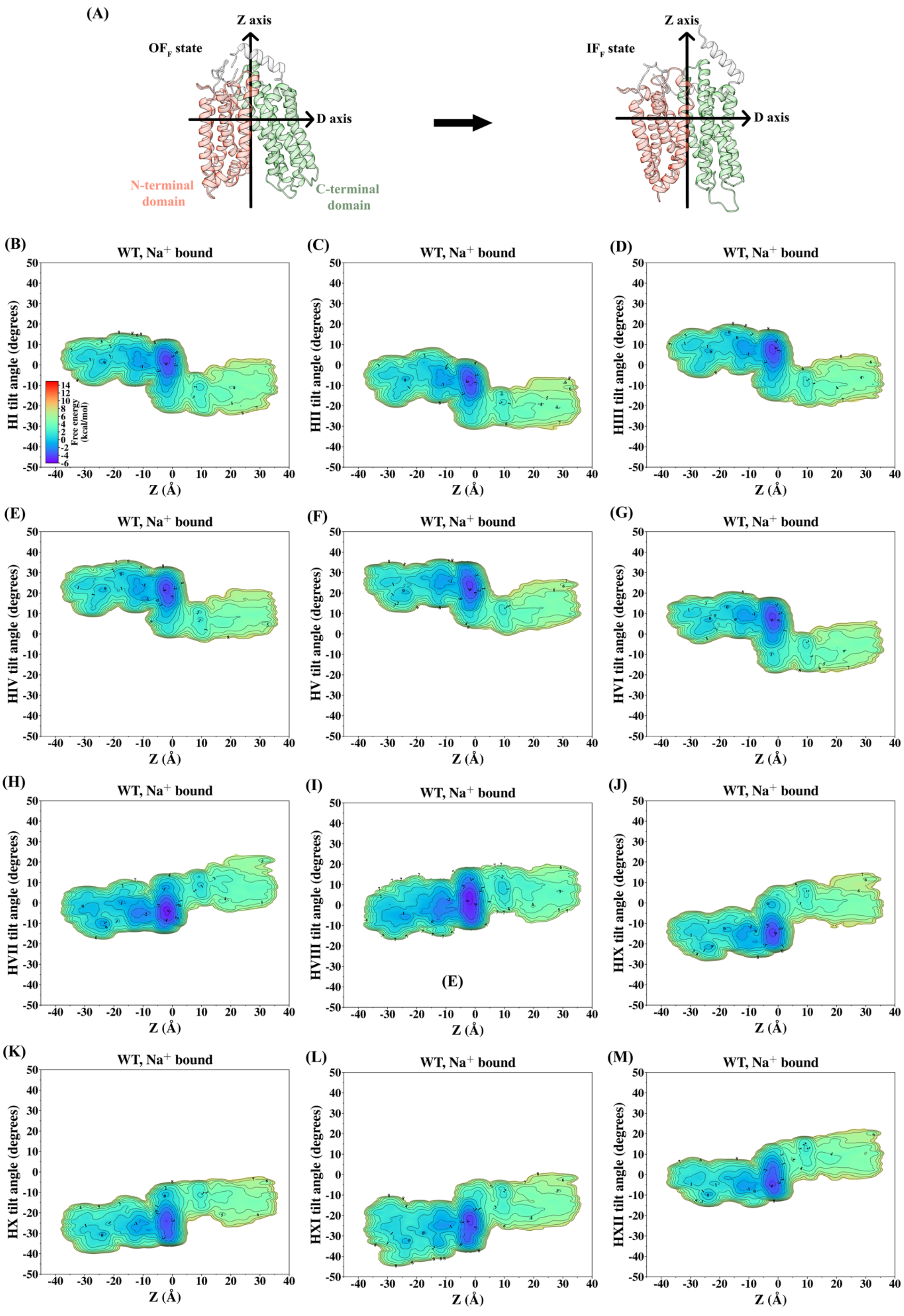

(A)
Z axis
OF_F state
D axis
N-terminal domain
C-terminal domain
Z axis
IF_F state
D axis
(B)
WT, Na+ bound
HI tilt angle (degrees)
Free energy (kcal/mol)
Z (Å)
(C)
WT, Na+ bound
HII tilt angle (degrees)
Z (Å)
(D)
WT, Na+ bound
HIII tilt angle (degrees)
Z (Å)
(E)
WT, Na+ bound
HIV tilt angle (degrees)
Z (Å)
(F)
WT, Na+ bound
HV tilt angle (degrees)
Z (Å)
(G)
WT, Na+ bound
HVI tilt angle (degrees)
Z (Å)
(H)
WT, Na+ bound
HVII tilt angle (degrees)
Z (Å)
(I)
WT, Na+ bound
HVIII tilt angle (degrees)
(E)
Z (Å)
(J)
WT, Na+ bound
HIX tilt angle (degrees)
Z (Å)
(K)
WT, Na+ bound
HX tilt angle (degrees)
Z (Å)
(L)
WT, Na+ bound
HXI tilt angle (degrees)
Z (Å)
(M)
WT, Na+ bound
HXII tilt angle (degrees)
Z (Å)

**Figure S4.** Free energy surfaces for the directional tilt angles of the transmembrane helices relative to the membrane normal during the melibiose translocation through WT MelB$_{St}$ in the $Na^{+}$ bound state. (A) Illustrative representation of the reorientation of the N-terminal (red) and C-terminal (green) transmembrane domains relative to the Z and X axes following the inward transport of melibiose (see Method for definition of axes and directional tilt angles). (B-G) N-terminal domain helices I to VI. (H-M), C-terminal domain helices VII to XII.

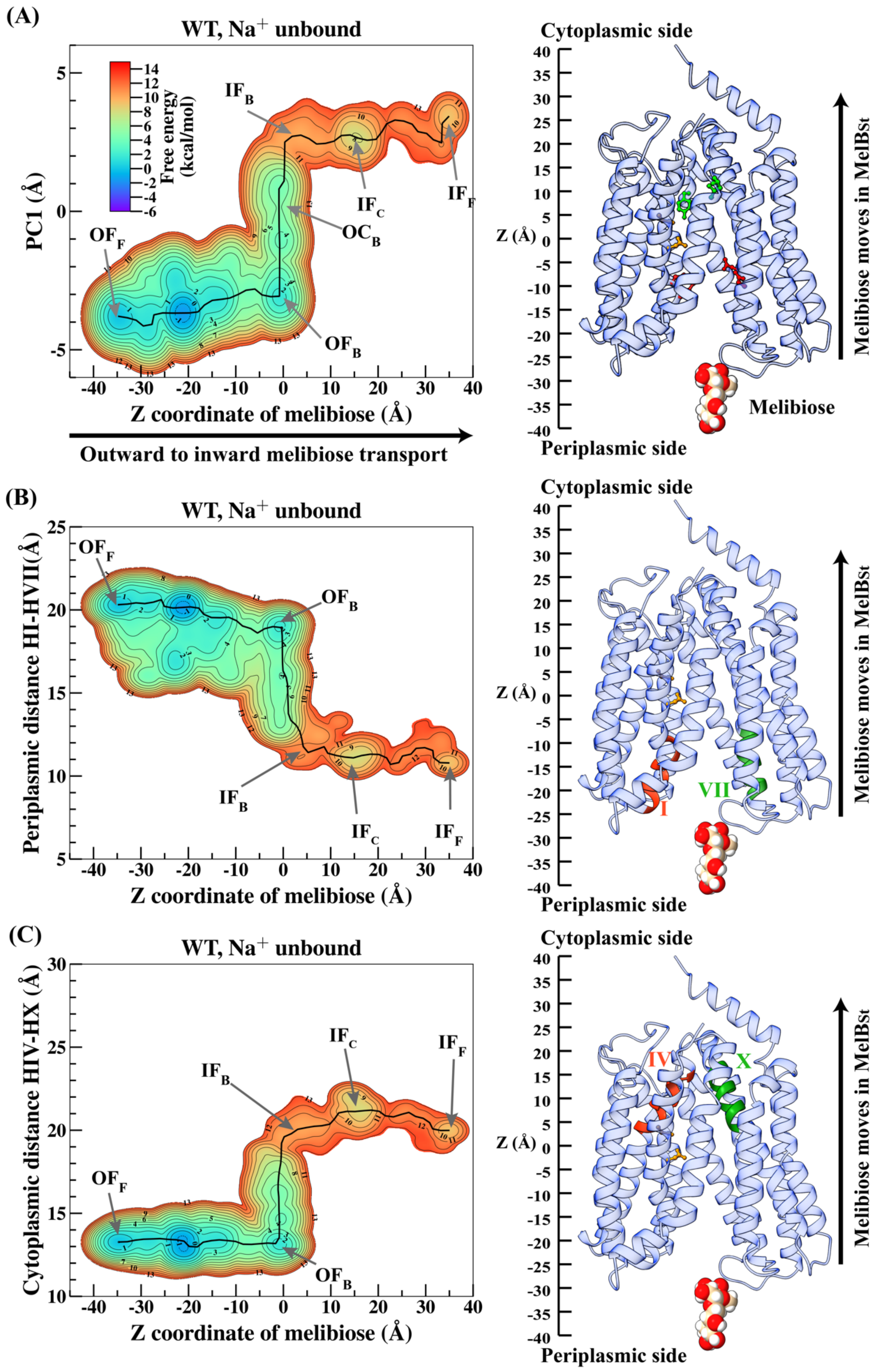

(A)
WT, Na+ unbound
Free energy (kcal/mol)
PC1 (Å)
Z coordinate of melibiose (Å)
Outward to inward melibiose transport
OF_F
OF_B
OC_B
IF_B
IF_C
IF_F
Cytoplasmic side
Periplasmic side
Z (Å)
Melibiose
Melibiose moves in MelBSt
(B)
WT, Na+ unbound
Periplasmic distance HI-HVII(Å)
Z coordinate of melibiose (Å)
OF_F
OF_B
IF_B
IF_C
IF_F
Cytoplasmic side
Periplasmic side
Z (Å)
I
VII
Melibiose moves in MelBSt
(C)
WT, Na+ unbound
Cytoplasmic distance HIV-HX (Å)
Z coordinate of melibiose (Å)
OF_F
OF_B
IF_B
IF_C
IF_F
Cytoplasmic side
Periplasmic side
Z (Å)
IV
X
Melibiose moves in MelBSt

**Figure S5.** Free energy surfaces (FES) for the transport of the melibiose through the WT $MelB_{St}$ in the $Na^+$ unbound state. (A) FES spanned by the first principle component of the backbone (PC1) vs. melibiose transport coordinate Z. An illustrative structure indicating the scale of the Z coordinate is shown on the right, and key residues in the periplasmic gate, binding site and cytoplasmic gate are highlighted in red, orange and green, respectively (see main text). (B) FES spanned by interhelical distance between helices I and VII on the periplasmic side. An illustrative structure indicating the helices I (red) and VII (green) is shown on the right. (C) FES spanned by interhelical distance between helices IV and X on the cytoplasmic side. An illustrative structure indicating the helices IV (red) and X (green) is shown on the right. The approximate minimum free energy pathways (MFEP) tracking the major basins on the FES are indicated as black lines, and the regions corresponding to the intermediate states are labeled by arrows. All FES plots share the same color bar as (A).

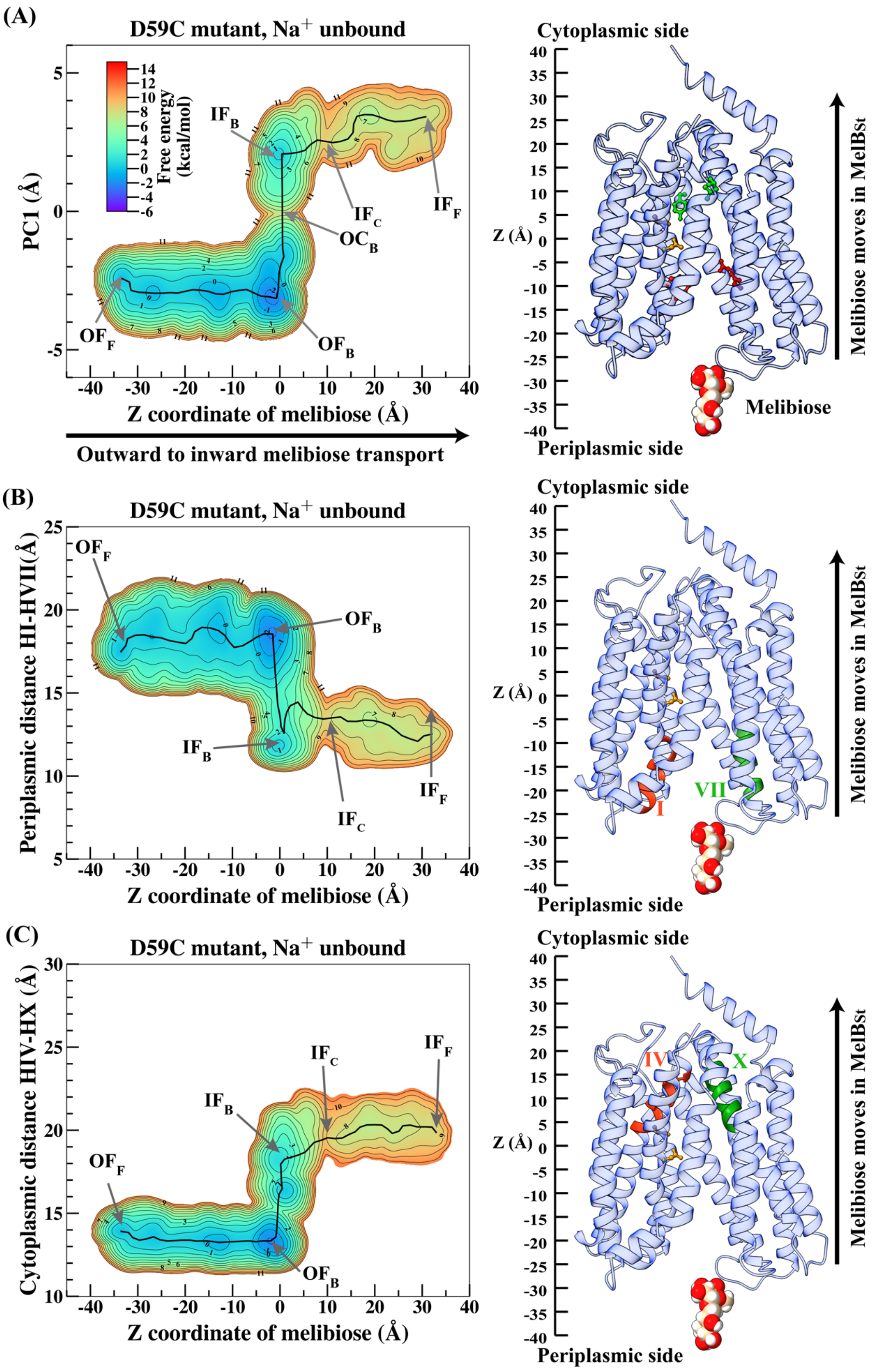

(A)
D59C mutant, Na$^+$ unbound
Free energy (kcal/mol)
PC1 (Å)
IF$_B$
IF$_C$
IF$_F$
OC$_B$
OF$_F$
OF$_B$
Z coordinate of melibiose (Å)
Outward to inward melibiose transport
Cytoplasmic side
Z (Å)
Melibiose moves in MelBSt
Melibiose
Periplasmic side
(B)
D59C mutant, Na$^+$ unbound
Periplasmic distance HI-HVII(Å)
OF$_F$
OF$_B$
IF$_B$
IF$_C$
IF$_F$
Z coordinate of melibiose (Å)
Cytoplasmic side
Z (Å)
I
VII
Melibiose moves in MelBSt
Periplasmic side
(C)
D59C mutant, Na$^+$ unbound
Cytoplasmic distance HIV-HX (Å)
IF$_C$
IF$_F$
IF$_B$
OF$_F$
OF$_B$
Z coordinate of melibiose (Å)
Cytoplasmic side
Z (Å)
IV
X
Melibiose moves in MelBSt
Periplasmic side

**Figure S6.** Free energy surfaces (FES) for the transport of the melibiose through the D59C mutant of $MelB_{St}$ in the $Na^+$ unbound state. (A) FES spanned by the first principle component of the backbone (PC1) vs. melibiose transport coordinate Z. An illustrative structure indicating the scale of the Z coordinate is shown on the right, and key residues in the periplasmic gate, binding site and cytoplasmic gate are highlighted in red, orange and green, respectively (see main text). (B) FES spanned by interhelical distance between helices I and VII on the periplasmic side. An illustrative structure indicating the helices I (red) and VII (green) is shown on the right. (C) FES spanned by interhelical distance between helices IV and X on the cytoplasmic side. An illustrative structure indicating the helices IV (red) and X (green) is shown on the right. The approximate minimum free energy pathways (MFEP) tracking the major basins on the FES are indicated as black lines, and the regions corresponding to the intermediate states are labeled by arrows. All FES plots share the same color bar as (A).

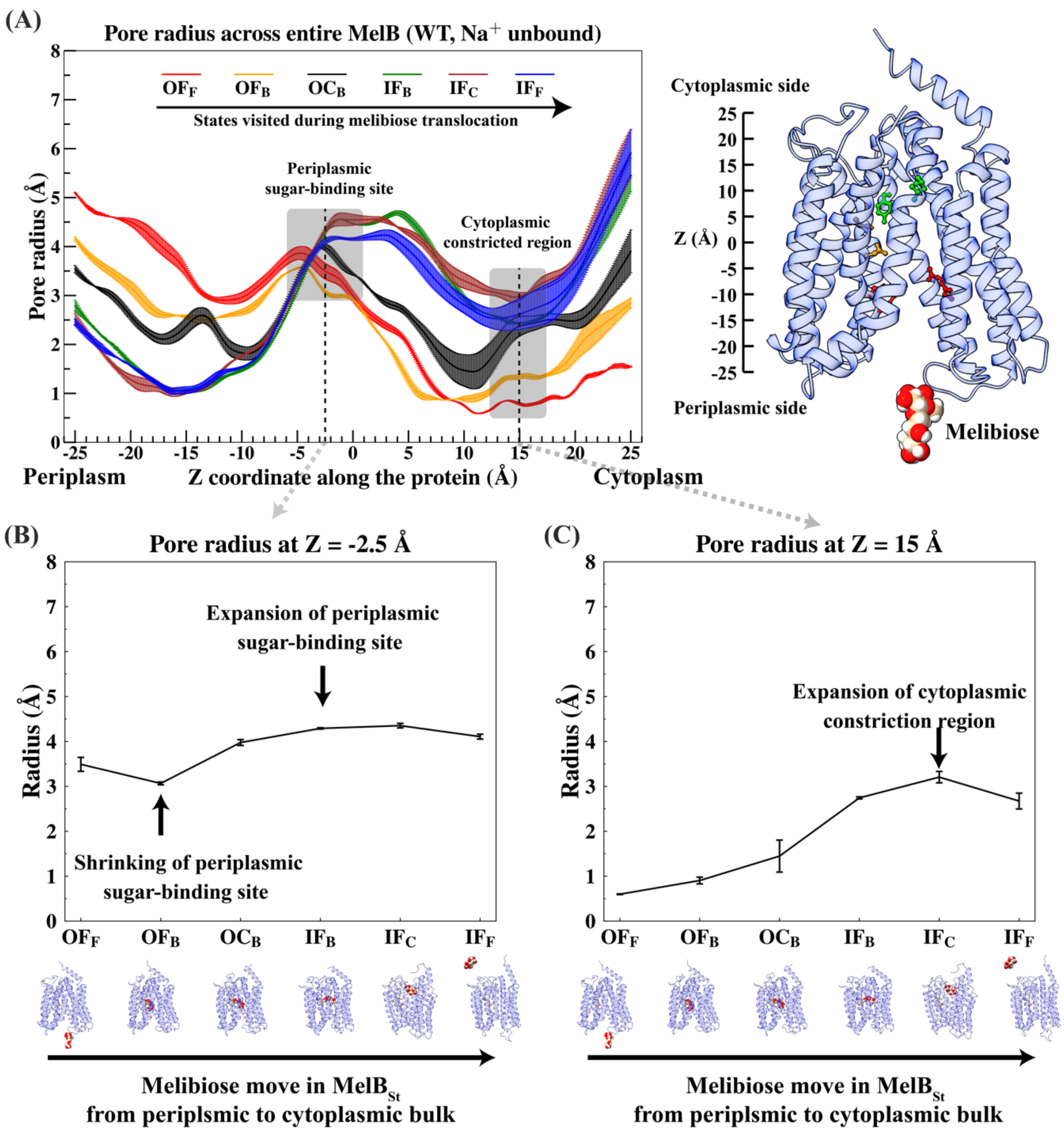


**Figure S7.** Change of pore radius profile coupled to melibiose transport through WT MelB$_{St}$ in the Na$^+$ unbound state. (A): Pore radii profile across the MelB as a function of the Z coordinate (relative to the COM of Asp124 and Asp19) along the entire protein (x-axis) for each different state during the melibiose transport process. As the melibiose is translocated from the periplasmic to cytoplasmic sides of the membrane, the system transitions from the OF$_F$ (red) to IF$_F$ (blue) states through the OF$_B$ (orange), OC$_B$ (black), IF$_B$ (green) and IF$_C$ (brown) intermediate states. The change in the states is coupled with gradual changes in the pore radius profile across the entire protein. The IF$_C$ state corresponds to the melibiose passing through the cytoplasmic constricted region near Z=15 Å. An illustrative structure indicating the scale of the Z coordinate is shown on the right. (B) and (C): Translocation of melibiose from the periplasmic to cytoplasmic sides of

$MelB_{St}$ induces the change of pore radii measured at Z=-2.5 Å (periplasmic sugar-binding site) and Z=15 Å (cytoplasmic constricted region). The melibiose translocation process is represented as the transitioning of the system from the $OF_F$ to $IF_F$ states through multiple intermediate states along the x-axes. The pore radii values in (B) and (C) are obtained from (A) using the crossing points between the pore radius profiles of each state with the vertical dashed lines at Z=-2.5 Å and Z=15 Å, respectively. The protein structures below the plots serve as visual guides for the location of the melibiose in different states.

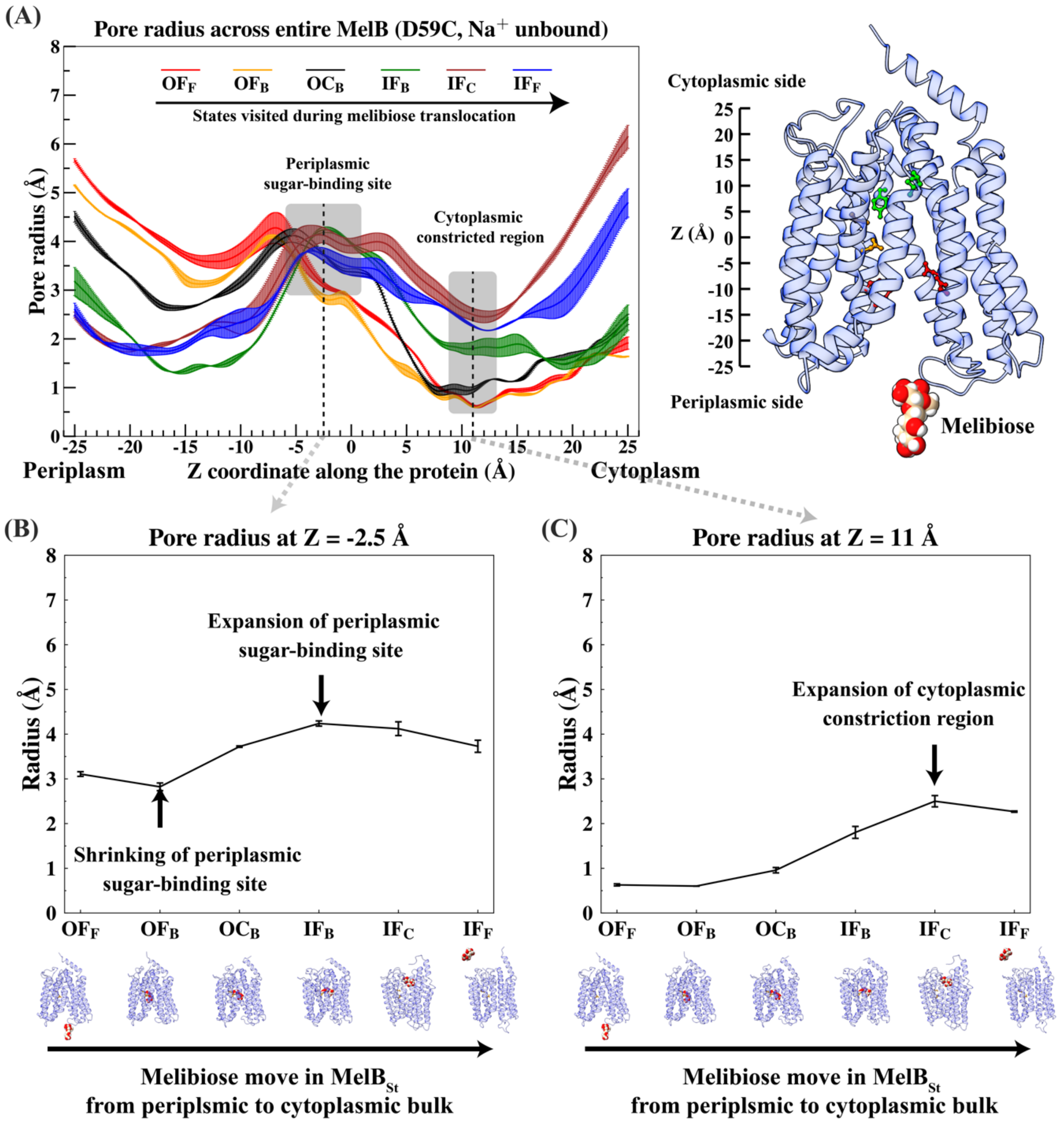


**Figure S8.** Change of pore radius profile coupled to melibiose transport through the D59C mutant of $MelB_{St}$ in the $Na^+$ unbound state. (A): Pore radii profile across the MelB as a function of the Z coordinate (relative to the COM of Asp124 and Asp19) along the entire protein (x-axis) for each different state during the melibiose transport process. As the melibiose is translocated from the periplasmic to cytoplasmic sides of the membrane, the system transitions from the $OF_F$ (red) to $IF_F$ (blue) states through the $OF_B$ (orange), $OC_B$ (black), $IF_B$ (green) and $IF_C$ (brown) intermediate states. The change in the states is coupled with gradual changes in the pore radius profile across the entire protein. The $IF_C$ state corresponds to the melibiose passing through the cytoplasmic constricted region near Z=11 Å. An illustrative structure indicating the scale of the Z coordinate is shown on the right. (B) and (C): Translocation of melibiose from the periplasmic to cytoplasmic sides of $MelB_{St}$ induces the change of pore radii measured at Z=-2.5 Å (periplasmic sugar-binding site) and Z=11 Å (cytoplasmic constricted region). The melibiose translocation process is represented as the transitioning of the system from the $OF_F$ to $IF_F$ states through multiple intermediate states along the x-axes. The pore radii values in (B) and (C) are obtained from (A) using the crossing points between the pore radius profiles of each state with the vertical dashed lines at Z=-2.5 Å and Z=11 Å, respectively. The protein structures below the plots serve as visual guides for the location of the melibiose in different states.

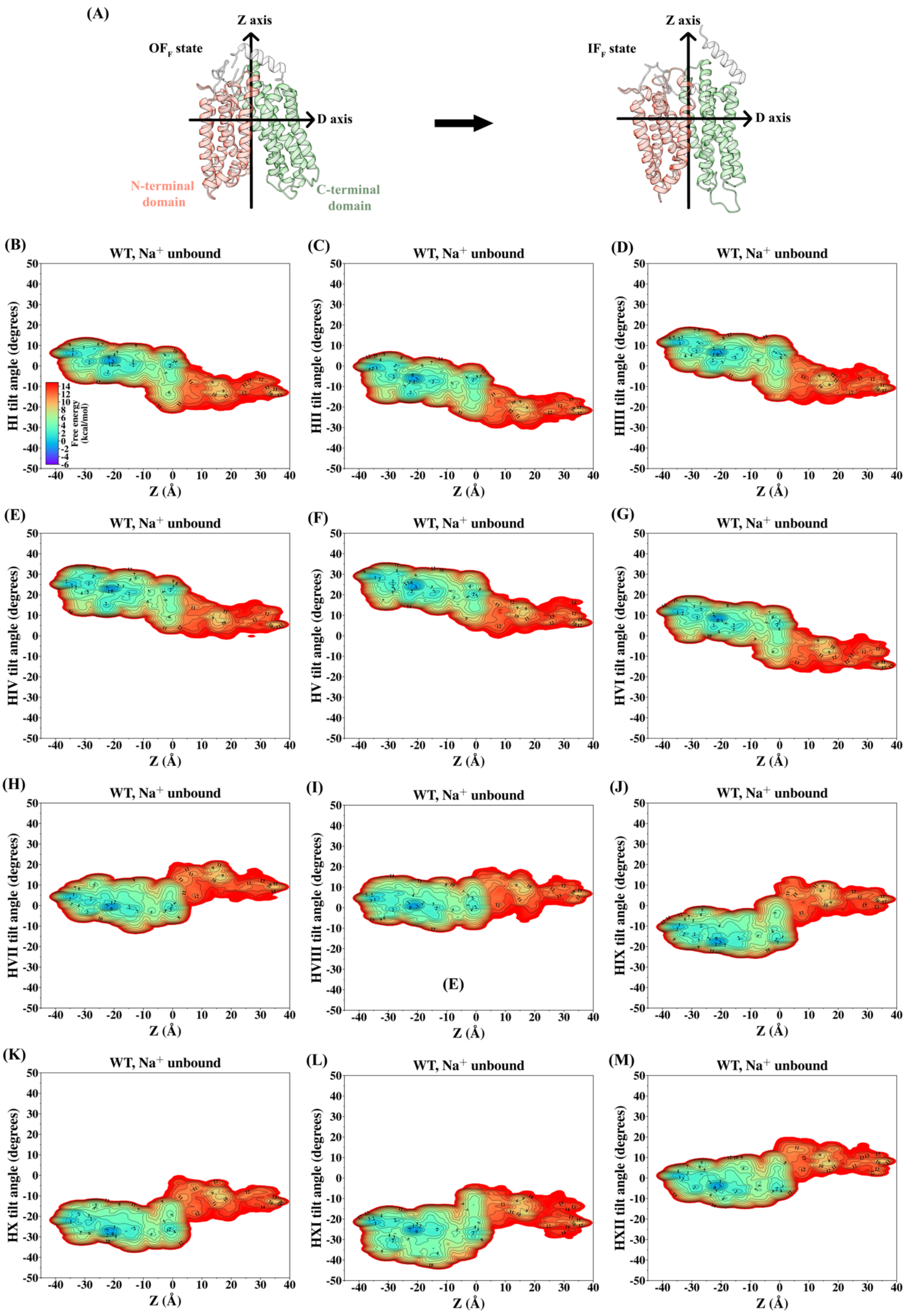
(A)
Z axis
OF_F state
D axis
N-terminal domain
C-terminal domain
Z axis
IF_F state
D axis
Free energy (kcal/mol)
(B)
WT, Na+ unbound
HI tilt angle (degrees)
Z (Å)
(C)
WT, Na+ unbound
HII tilt angle (degrees)
Z (Å)
(D)
WT, Na+ unbound
HIII tilt angle (degrees)
Z (Å)
(E)
WT, Na+ unbound
HIV tilt angle (degrees)
Z (Å)
(F)
WT, Na+ unbound
HV tilt angle (degrees)
Z (Å)
(G)
WT, Na+ unbound
HVI tilt angle (degrees)
Z (Å)
(H)
WT, Na+ unbound
HVII tilt angle (degrees)
Z (Å)
(I)
WT, Na+ unbound
HVIII tilt angle (degrees)
(E)
Z (Å)
(J)
WT, Na+ unbound
HIX tilt angle (degrees)
Z (Å)
(K)
WT, Na+ unbound
HX tilt angle (degrees)
Z (Å)
(L)
WT, Na+ unbound
HXI tilt angle (degrees)
Z (Å)
(M)
WT, Na+ unbound
HXII tilt angle (degrees)
Z (Å)

**Figure S9.** Free energy surfaces for the directional tilt angles of the transmembrane helices relative to the membrane normal during the melibiose translocation through WT $MelB_{St}$ in the $Na^+$ unbound state. (A) Illustrative representation of the reorientation of the N-terminal (red) and C-terminal (green) transmembrane domains relative to the Z and X axes following the inward transport of melibiose (see Method for definition of axes and directional tilt angles). (B-G) N-terminal domain helices I to VI. (H-M), C-terminal domain helices VII to XII.

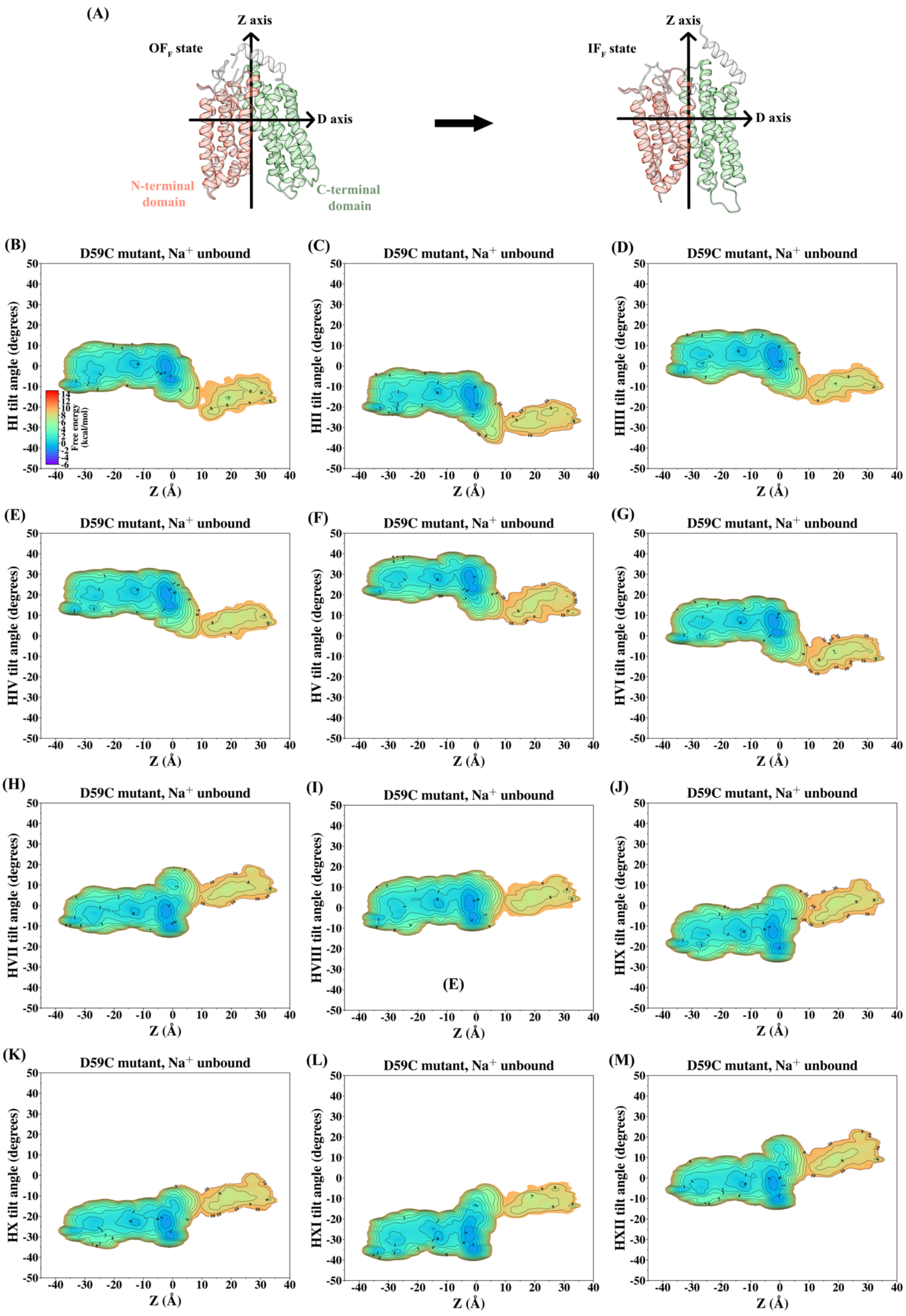

(A)
Z axis
OF$_F$ state
D axis
N-terminal domain
C-terminal domain
IF$_F$ state
(B)
D59C mutant, Na$^+$ unbound
HI tilt angle (degrees)
Z (Å)
Free energy (kcal/mol)
(C)
D59C mutant, Na$^+$ unbound
HII tilt angle (degrees)
Z (Å)
(D)
D59C mutant, Na$^+$ unbound
HIII tilt angle (degrees)
Z (Å)
(E)
D59C mutant, Na$^+$ unbound
HIV tilt angle (degrees)
Z (Å)
(F)
D59C mutant, Na$^+$ unbound
HV tilt angle (degrees)
Z (Å)
(G)
D59C mutant, Na$^+$ unbound
HVI tilt angle (degrees)
Z (Å)
(H)
D59C mutant, Na$^+$ unbound
HVII tilt angle (degrees)
Z (Å)
(I)
D59C mutant, Na$^+$ unbound
HVIII tilt angle (degrees)
(E)
Z (Å)
(J)
D59C mutant, Na$^+$ unbound
HIX tilt angle (degrees)
Z (Å)
(K)
D59C mutant, Na$^+$ unbound
HX tilt angle (degrees)
Z (Å)
(L)
D59C mutant, Na$^+$ unbound
HXI tilt angle (degrees)
Z (Å)
(M)
D59C mutant, Na$^+$ unbound
HXII tilt angle (degrees)
Z (Å)

**Figure S10.** Free energy surfaces for the directional tilt angles of the transmembrane helices relative to the membrane normal during the melibiose translocation through the D59C mutant of WT $MelB_{St}$ in the $Na^+$ unbound state. (A) Illustrative representation of the reorientation of the N-terminal (red) and C-terminal (green) transmembrane domains relative to the Z and X axes following the inward transport of melibiose (see Method for definition of axes and directional tilt angles). (B-G) N-terminal domain helices I to VI. (H-M), C-terminal domain helices VII to XII.

(A)

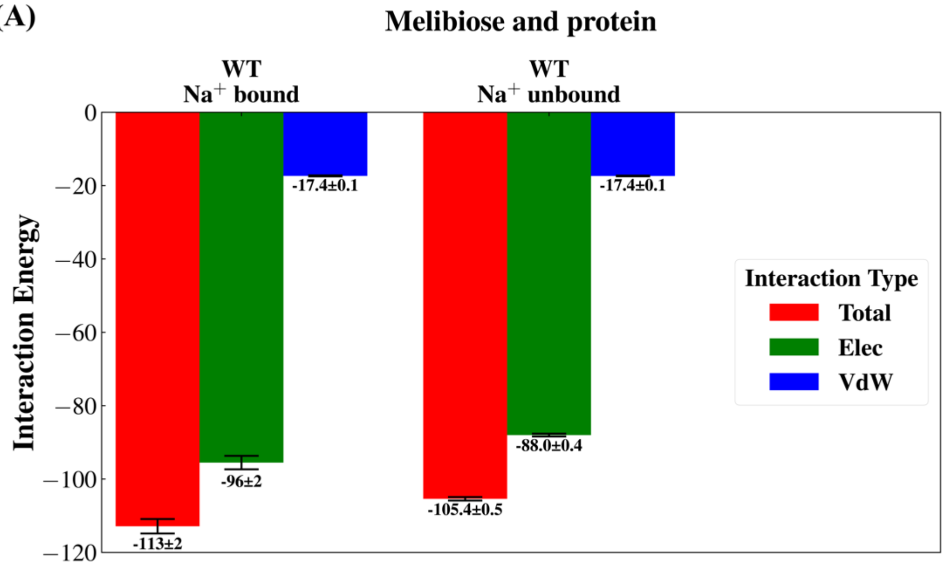


(B)

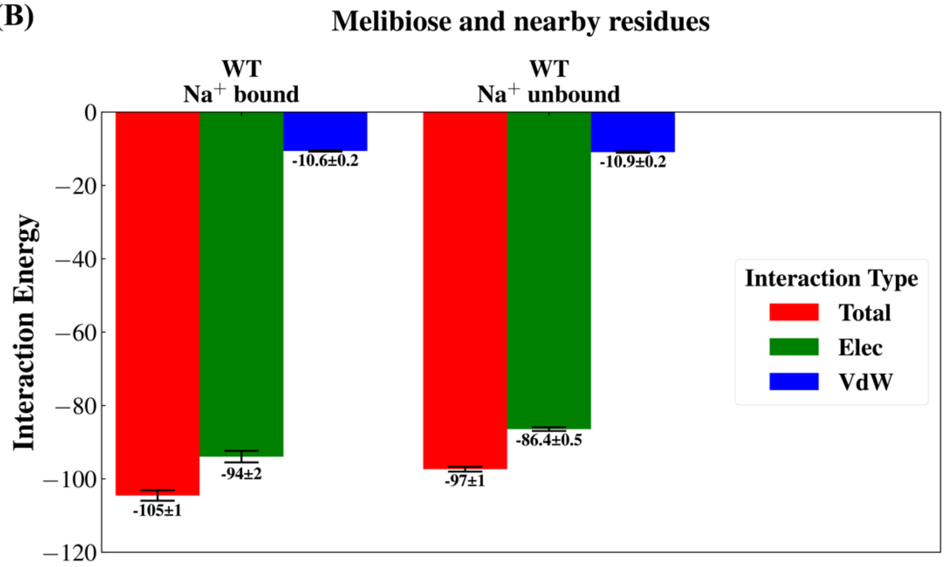


**Figure S11.** Comparison of the protein-melibiose interaction energy in the $OF_B$ state of the WT $Na^+$ bound, WT $Na^+$ unbound systems. (A) The average interaction energies between the bound melibiose and the whole protein in the $OF_B$ state. (B) The average interaction energy between the bound melibiose and its nearby residues (Lys18, Asp19, Tyr26, Arg52, Tyr120, Asp124, Trp128, Arg149, Trp342, Thr373, and Val376). The total non-bonded interaction energies (red) are decomposed into electrostatic (blue) and van der Waals (blue) interaction energies. For each system, all interaction energies are calculated using a total of 500 ns unbiased trajectory sampling the $OF_B$ state. Error bars are estimated by block-average analysis.

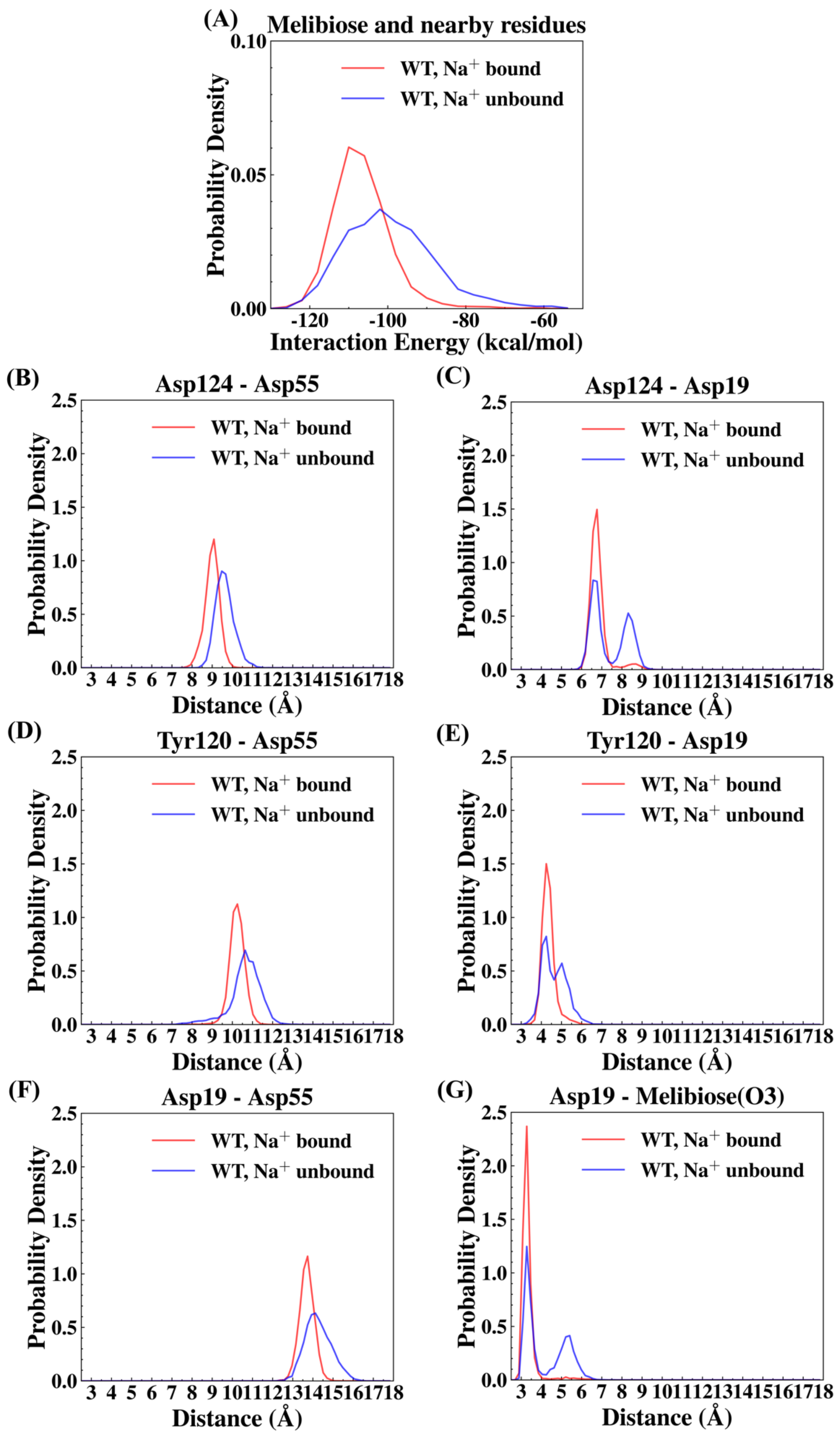

(A)
Melibiose and nearby residues
WT, Na+ bound
WT, Na+ unbound
Probability Density
Interaction Energy (kcal/mol)
(B)
Asp124 - Asp55
(C)
Asp124 - Asp19
(D)
Tyr120 - Asp55
(E)
Tyr120 - Asp19
(F)
Asp19 - Asp55
(G)
Asp19 - Melibiose(O3)
Distance (Å)

**Figure S12.** $Na^+$ unbinding from the cation-binding site alters the melibiose-protein interactions at the melibiose-binding site. The motions of Asp124 and Tyr120 are involved in the allosteric coupling between the cation- and sugar-binding sites. The Asp55 and Asp19 represent key residues in the cation- and sugar-binding sites, respectively. (A) Probability distribution of the interaction energy between the melibiose and nearby residues (defined in Fig. 8 and main text) in the WT $Na^+$ bound, WT $Na^+$ unbound systems. (B-D) Probability distributions of the distances between the terminal carboxyl carbon atoms on the side chains of (B) Asp55 and Asp19 (C) Asp124 and Asp55 (D) Asp124 and Asp19 in the two systems. (E-F) Probability distributions of the distances between the terminal hydroxyl oxygen atom on the side chain of Tyr120 and the terminal carboxyl carbon atom of (E) Asp55 (F)Asp19 in the two systems. (G) Probability distribution of the distance between the terminal carboxyl carbon atom of Asp19 and the oxygen atom of the 3-hydroxyl group of the melibiose. $Na^+$ unbinding from the cation-binding site increases all of the above-mentioned distances, leading to structural perturbations at the sugar-binding site such as a loosened Asp19-melibiose hydrogen bond. For each system, all distributions are calculated using a total of 500 ns unbiased trajectory sampling the $OF_B$ state.

**Table S1.** The average number of hydrogen bonds between melibiose and protein in the $OF_B$ states. The average numbers of hydrogen bonds between melibiose and key individual residues at the binding site (Asp19, Asp124, Arg149 and Trp128) are also listed.

| Type of hydrogen bonds | System | |
|---|---|---|
| | WT, $Na^+$ bound | WT, $Na^+$ unbound |
| Melibiose-Protein (total) | 6.2 ± 0.1 | 5.3 ± 0.2 |
| Melibiose-Asp19 | 0.9 ± 0.1 | 0.5 ± 0.1 |
| Melibiose-Asp124 | 1.79 ± 0.01 | 1.6 ± 0.1 |
| Melibiose-Arg149 | 2.53 ± 0.03 | 2.3 ± 0.2 |
| Melibiose-Trp128 | 0.68 ± 0.02 | 0.6 ± 0.1 |
| Melibiose-Lys18 | 0.2 ± 0.1 | 0.09 ± 0.04 |
